# TUTase mediated site-directed access to clickable chromatin employing CRISPR-dCas9

**DOI:** 10.1101/846980

**Authors:** Jerrin Thomas George, Mohd. Azhar, Meghali Aich, Dipanjali Sinha, Uddhav B. Ambi, Souvik Maiti, Debojyoti Chakraborty, Seergazhi G. Srivatsan

**Affiliations:** Department of Chemistry, Indian Institute of Science Education and Research (IISER), Pune Dr. Homi Bhabha Road, Pune 411008, India; Genomics and Molecular Medicine Unit, Council of Scientific and Industrial Research-Institute of Genomics & Integrative Biology, New Delhi, 110025, India; Institute of Genomics and Integrative Biology (IGIB)-National Chemical Laboratory (NCL) Joint Center, Council of Scientific and Industrial Research–National Chemical Laboratory, Pune, 411008, India

## Abstract

Locus-specific interrogation of the genome using programmable CRISPR-based technologies is tremendously useful in dissecting the molecular basis of target gene function and modulating its downstream output. Although these tools are widely utilized in recruiting genetically encoded functional proteins, display of small molecules using this technique is not well developed due to inadequate labeling technologies. Here, we report the development of a modular technology, sgRNA-Click (sgR-CLK), which harnesses the power of bioorthogonal click chemistry for remodeling CRISPR to display synthetic molecules on target genes. A terminal uridylyl transferase (TUTase) was repurposed to construct an sgRNA containing multiple minimally invasive bioorthogonal clickable handles, which served as a Trojan horse on CRISPR-dCas9 system to guide synthetic tags site-specifically on chromatin employing copper-catalyzed or strain-promoted click reactions. Our results demonstrate that sgR-CLK could provide a simplified solution for site-directed display of small molecules to study as well as modulate the function of gene targets.

## Introduction

The bacterial adaptive defense system, clustered regularly interspaced short palindromic repeats (CRISPR)–CRISPR-associated proteins (Cas), is in spotlight for its ability to serve as a powerful genome targeting and editing tool^1,2^. The ease-of-use and flexibility to target any genomic loci by simply changing the sequence of the single guide RNA (sgRNA) sets this technology apart from other traditionally used technologies like zinc finger nucleases (ZFNs) and transcription activator-like effector nucleases (TALENs), which requires extensive protein engineering to target specific DNA bases^3–5^. The catalytically inactive mutant of Cas9, also called dead Cas9 (dCas9), has further harnessed the utility of this genome targeting tool for localized delivery of functional tags to enable transcriptional activation/repression^6,7^, base-editing^8,9^, chromatin pull-down^10,11^, and gene visualization^12^. These applications greatly rely on dCas9 derivatized with genetically encoded fusion proteins (e.g., eGFP, APOBEC, VP64)^13^ and epitope tags (e.g., Flag and SunTag)^14–16^. Similarly, engineered sgRNA scaffolds with recruitment sequences for RNA-binding proteins or aptamers have also been used to regulate as well as visualize specific genes^17–19^ While tools based on dCas9-sgRNA have been widely utilized in delivering functional proteins, locus-specific delivery of synthetic molecules such as drugs, photoactivatable groups, pull-down probes, and epigenomic and transcription modulators remains a technical challenge due to the lack of adequate labeling technologies^20^. Therefore, development of practical labeling technologies to recruit such small molecule probes and effectors will not only allow the profiling/modulation of the interaction partners of the target gene but also would profoundly expand the therapeutic and diagnostic potential of CRISPR system^21–23^.

Towards this direction, methods including maleimide chemistry, post-synthetic modification of a HaloTag and incorporation of unnatural amino acids have been used to derivatize dCas9^24,25^. These ligation reactions are not per se bioorthogonal and, hence could result in high background due to non-specific reactions in cellular environment. More recently, expression platforms composed of peptide-conjugated dCas9 and ligation modules have been used to (i) introduce small molecule epigenetic baits using split intein^20^ and (ii) transfer biotin using biotin ligase for chromatin capture^26^. However, engineering dCas9 with small molecules using expression platforms is complex, and further, the efficiency of conjugation is dependent on the functional moiety tagged on the ligation module^27^. On the other hand, covalent derivatization of sgRNA is envisioned as a powerful alternative to localize small molecule functional tags to gene loci^28,29^. For example, sgRNA tagged with a fluorescent dye, generated by splint-ligating a chemically synthesized RNA oligonucleotide, was used in establishing a CRISPR-based FISH assay to visualize gene loci in fixed cells^24^. However, except for a very few examples, accessing long and structured RNAs like sgRNA labeled with a wide range of functional tags remains a major challenge due to known drawbacks in conventional chemical and enzymatic labeling approaches. Further, dCas9 and sgRNA conjugation strategies rely on functional tagging prior to the formation of the active ternary complex (dCas9:sgRNA:target DNA)^18^. Hence, attaching synthetic modulators/probes in this way could potentially perturb the ribonucleoprotein (RNP) complex formation or its binding efficiency to the target DNA sequence^28^. In this regard, we hypothesized that bioorthogonal click reaction would serve as a powerful tool in installing noninvasive reactive handles on sgRNA, which could be further functionalized chemo-selectively upon formation of the ternary complex for downstream applications^30,31^.

Here, we describe the development of an innovative labeling technology to display minimally perturbing clickable groups (e.g., azide) at a genetic locus of interest by repurposing terminal uridylyl transferase (TUTase) to incorporate azide-modified UTP analogs site-specifically at the 3’ end of sgRNAs. An azide-tailed sgRNA constructed using TUTase served as a Trojan horse on CRISPR-dCas9 system to direct functional tags on telomeres. Indeed, *in situ* click reaction performed on the target-bound ternary complex using biotin-alkynes showed significant enrichment of the telomeric DNA repeat upon pull-down. This technique, which we named sgRNA-Click (sgR-CLK), is modular as any sgRNA of choice can be engineered to deploy desired synthetic modulators/probes, thereby offering new means to interrogate gene loci of interest.

## Results

### Blueprint of sgR-CLK

Our strategy to construct clickable sgRNA so as to display functional tags on a specific gene target using CRISPR system is based the following consideration. Among various clickable groups, azide is structurally minimally perturbing and can be used in different bioorthogonal reactions like copper-catalyzed azide-alkyne cycloaddition (CuAAC), strain-promoted azide-alkyne cycloaddition (SPAAC) and Staudinger ligation reactions to install variety of functionalities on biomolecules^32,33^. However, chemical labeling of RNA with azide groups is not straightforward as azide-modified phosphoramidites are not stable under solid-phase oligonucleotide synthesis conditions^34^. Further, indiscriminate body labeling of sgRNA utilizing RNA polymerases is likely to perturb the formation of an active CRISPR RNP complex. To overcome this, we decided to repurpose the RNA tailing ability of a TUTase (SpCID1) in site-specifically derivatizing sgRNA of choice with multiple azide groups at the 3’ end employing azide-modified UTP analogs. An sgRNA thus obtained would enable the display of desired functional tags on the target gene by either post-hybridization (Figure 1, path A) or prehybridization click reaction (Figure 1, path B) with appropriate alkyne-labeled tags. In fact, our results demonstrated that post-hybridization click strategy is the method of choice for installing synthetic cargos.

**Figure 1.**
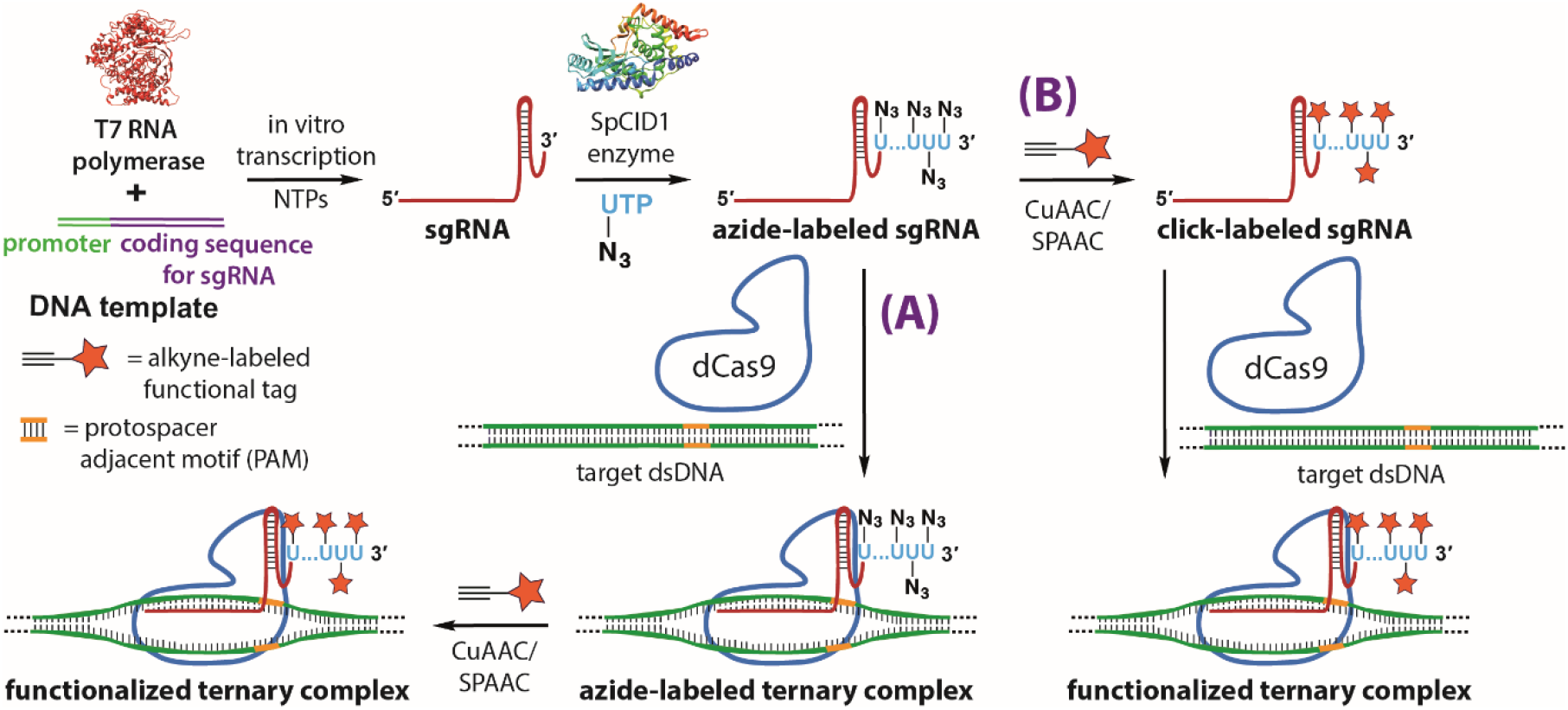
sgR-CLK design to display functional tags on a specific gene. *In vitro* transcribed CRISPR sgRNA is terminal uridylated with an azide-labeled nucleotide analog using SpCID1. Azide-labeled sgRNA can be further used to generate functionalized ternary complex by either post-hybridization (path **A**) or pre-hybridization (path **B**) click reaction with an alkyne counterpart containing the desired function tag.

### Azide-tailed sgRNAs using TUTase

The prerequisite to establish sgR-CLK would require TUTase to efficiently incorporate azide-modified UTP analogs into highly structured sgRNA sequences. TUTases play a key role in epitranscriptomics wherein these enzymes add multiple uridylates at the 3’ end of RNA, which signals RNA biogenesis, processing, degradation or stabilization^35,36^. In the family of TUTases, a cytoplasmic TUTase from *Schizosaccharomyces pombe* known as caffeine-induced death suppressor 1 (SpCID1) has been shown as an important posttranscriptional RNA labeling enzyme^37,38^. Careful examination of the crystal structure of SpCID1 in the presence of UTP revealed that the C5-position of uridine does not make contacts with neighboring amino acid residues (Figure 2a)^39^. This observation suggested that the enzyme might be promiscuous to C5-modified UTP analogs. Therefore, we synthesized azide-modified UTP analogs namely, AMUTP, APUTP and ATUTP with increasing linker length having a methyl, propyl and tetraethylene glycol spacer, respectively, between the azide group and nucleobase (Figure 2b)^40,41^. We then cloned SpCID1 gene from *S. pombe* fission yeast and expressed the recombinant protein in *E. coli* (see supplementary information for details, Figure S1-S4, Table S1). The efficacy of SpCID1 to incorporate the nucleotide analogs was evaluated by using a model RNA oligonucleotide sequence bearing a fluorescein (FAM) label at the 5’ end (Figure 2c and Table 1).

**Figure 2.**
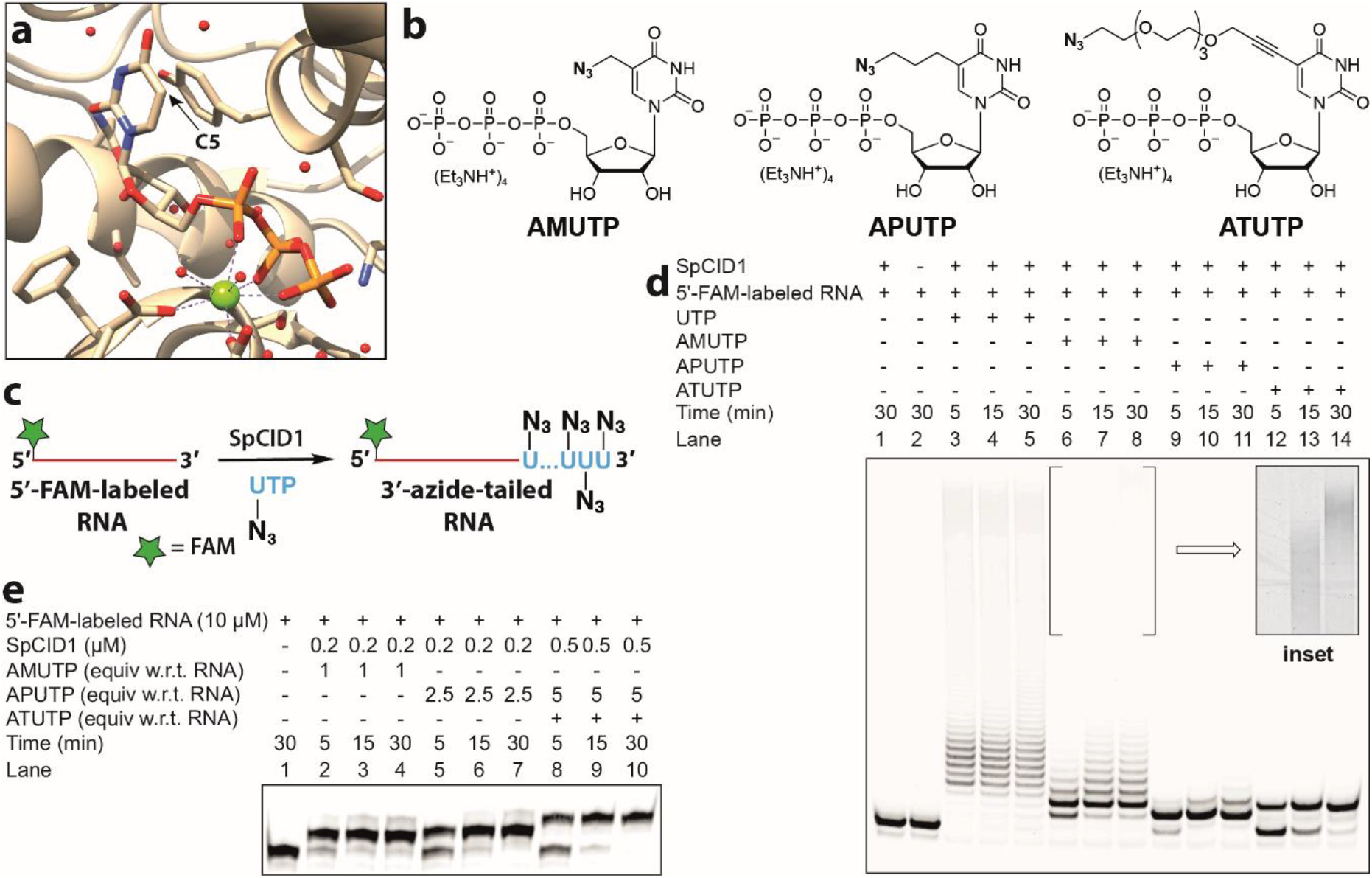
SpCID1, a TUTase enzyme, efficiently incorporates C5-azide-modified nucleotides at the 3’ end of an RNA oligonucleotide. **(a)** Crystal structure of SpCID1 bound to UTP (PDB 4FH5) generated using UCSF Chimera software^39^. C5 position of uridine triphosphate is shown using an arrow. **(b)** Structure of azide-modified UTP analogs used in terminal uridylation reaction. **(c)** Scheme showing 3’-terminal uridylation of model 5’-FAM-labeled RNA with modified UTP analogs. **(d)** Image of PAGE resolved reaction products of terminal uridylation reactions in the presence of UTP/modified UTPs using SpCID1. Inset: Processive incorporation of AMUTP was evident upon increasing the contrast of the gel image. **(e)** Gel image showing optimized reaction conditions to effect single nucleotide incorporation. Experiments **d** and **e** were performed in n = 2 independent experiments.

**Table 1.**
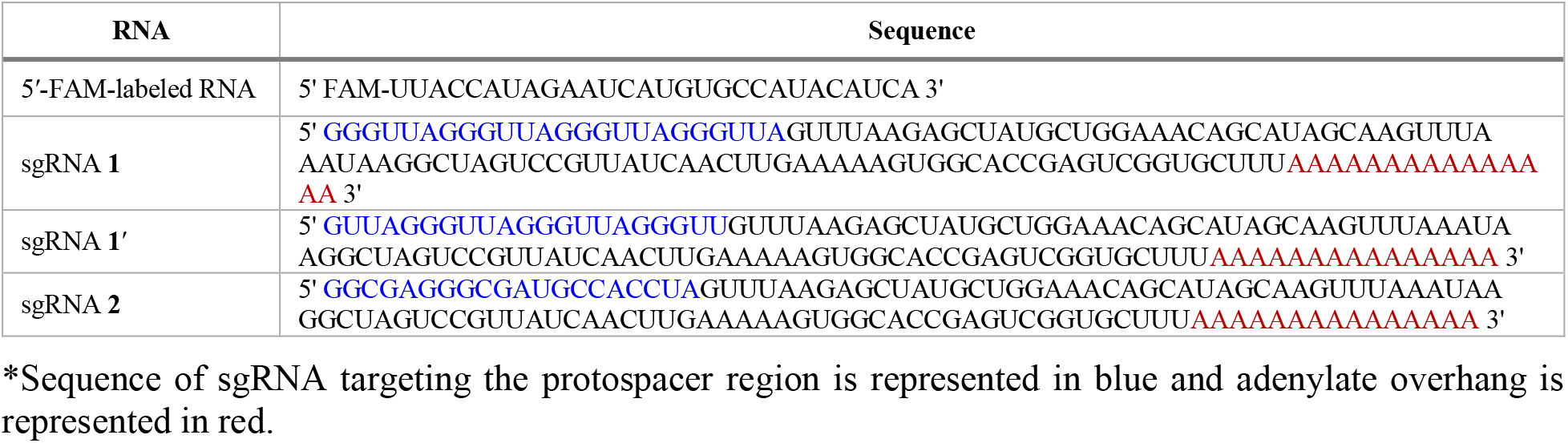
Sequence of model RNA oligonucleotide and sgRNAs.

SpCID1 was incubated with 5’-FAM-labeled RNA in the presence of UTP or azide-modified UTPs. Reaction aliquots at different time intervals were resolved by polyacrylamide gel electrophoresis (PAGE) under denaturing conditions and the reaction products were imaged using a fluorescence scanner (Figure 2d). A biphasic incorporation pattern corresponding to a distributive and processive addition of UTP, consistent with literature report, was observed (lane 3–5)^42^. Distributive enzymatic UMP addition involves the dissociation of enzyme from substrate RNA after every successive UMP addition giving rise to RNA oligonucleotides containing few added nucleotides (6–9 nucleotides). However, in the processive phase the enzyme adds several nucleotides to the tailed RNA substrates produced in the distributive phase, which is thought to be a result of increase in binding affinity of enzyme to uridylated RNA^42^. Remarkably, all the nucleotide analogs served as good substrates in the terminal uridylation reaction. Notably, the analogs were predominantly incorporated via distributive enzyme incorporation, wherein the number of additions dependent on the length of the linker. While AMU was incorporated multiple times (lane 6–8), APUTP and ATUTP produced RNA oligonucleotides containing one or two nucleotide analogs (lane 9–14). Further optimization by varying enzyme-nucleotide stoichiometry yielded singly-modified RNA oligonucleotide as the major product, which was characterized by mass analysis (Figure 2e, Figure S5, S6 and Table S2). AMU-labeled RNA oligonucleotide was readily functionalized by SPAAC and CuAAC reactions using Cy3-DBCO strained alkyne and Alexa 594-alkyne, respectively (Figure 3). It is worth mentioning here that RNA containing an azide label at the 3’ end can be used to generate RNA with terminally- or internally-labeled biophysical probes by click reaction or ligation followed by click reaction^43^. Recently, chemically synthesized crRNAs containing chemo-selective reactive handles have been used in stitching tracrRNA to yield full-length sgRNA or donor DNA to facilitate homology directed repair^44,45^.

**Figure 3.**
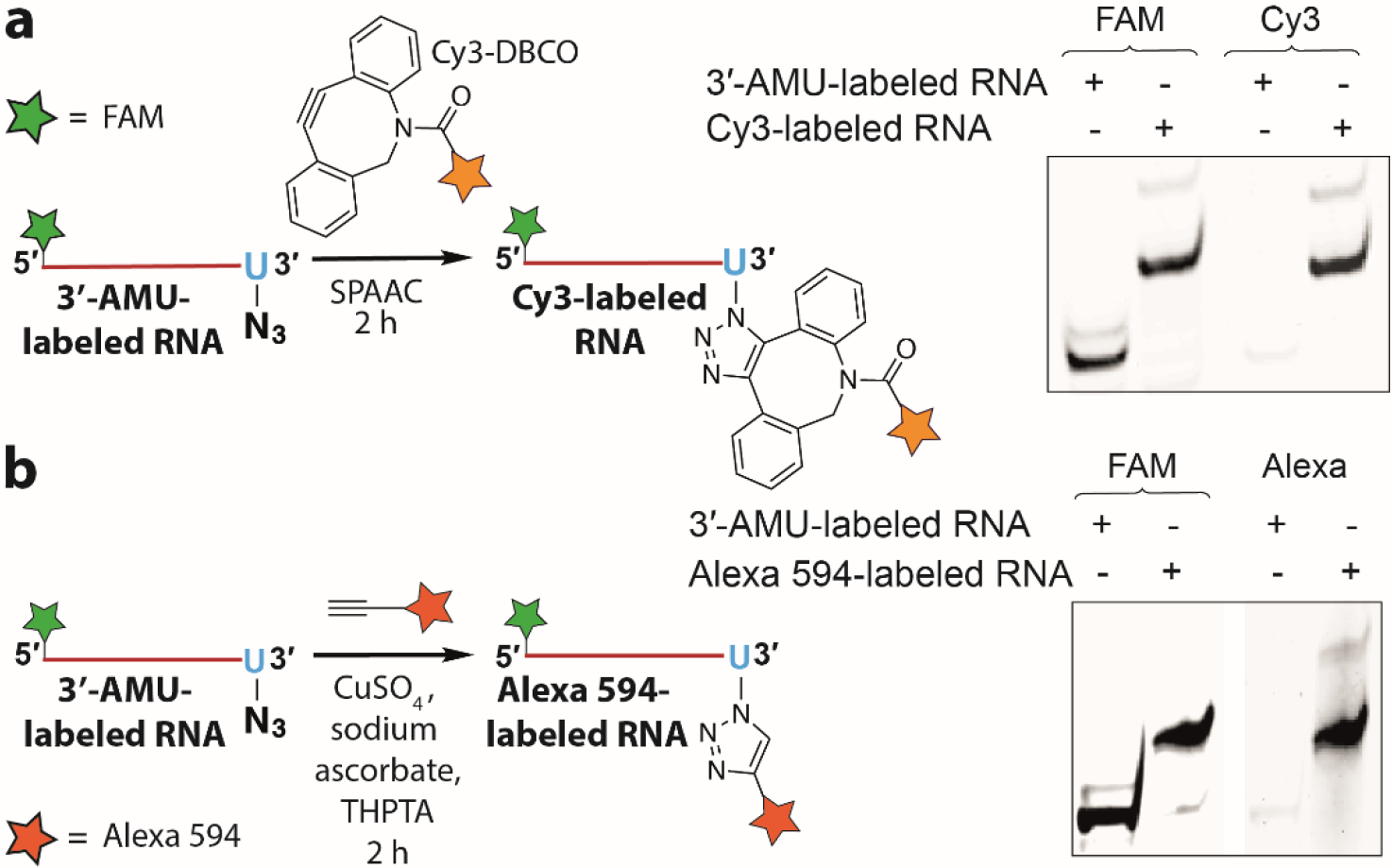
AMU-labeled RNA oligonucleotide is compatible for click functionalization. The gels were scanned at FAM and Cy3 wavelengths. (a) Scheme and gel image of SPAAC reaction between 3’-AMU-labeled RNA and Cy3-DBCO. (b) CuAAC reaction between 3’-AMU-labeled RNA and Alexa 594 alkyne. Structure of alkyne substrates is provided in Figure S7.

However, this strategy is not suitable for introducing functional tags on sgRNA unlike our method, which introduces multiple free azide labels on sgRNA for further functionalization. Hence, in the present study, we preferred to synthesize sgRNAs containing multiple AMU residues at its 3’ end utilizing its minimally perturbing nature, which in turn may not affect the formation of the active CRISPR complex for subsequent display of multiple functional tags by click chemistry.

As a study system to establish sgR-CLK, we designed sgRNAs with a protospacer targeting telomere repeat region and eGFP gene. We used telomere as the target since CRISPR-based tools developed to visualize and pull-down specific genes commonly use this sequence as a model. eGFP was used as a non-targeting sequence. We initially used a conventional guide RNA design^12^ to synthesize sgRNA targeting the telomere region by *in vitro* transcription reaction using T7 RNA polymerase and template CT1 (Table S1). Upon incubation with SpCID1 and AMUTP we observed no detectable incorporation of the nucleotide analog (data not shown). This is possibly because the trans-acting CRISPR RNA (tracrRNA) region of sgRNA being RNA pol III transcribed, ends with a strong hairpin loop structure at its 3’ end (Figure S8a)^46^. This feature is also observed in the crystal structure of guide RNA bound to Cas9 and target DNA^47^. *In vitro* experiments have revealed that SpCID1 requires a ~13 nucleotide stretch of single stranded RNA for efficient binding and uridylation^37^. Possibly due to this reason the enzyme failed to incorporate AMUTP into the strongly structured 3’ end of sgRNA.

To circumvent this problem, we took the cue from the abilty of SpCID1 to bind and uridylate unstructured poly(A) tail region of mRNA^38^. Additionally, poly(A) and poly(U) tracts at the 3’ end of guide RNA has been reported to increase the stability and efficiency of CRISPR systems^48,49^. Based on these key observations, we decided to recofigure sgRNA to contain a poly(A) tail at the 3’ end with the view that it would facilitate efficient incorporation of AMUTP. This notion was also supported by secondary structure prediction, which suggested that extension of sgRNA with a 15-mer homopolymer of adenosine preserved the overall structure of the guide RNA and introduced an unstructured 3’-overhang (Figure S8b).

PCR amplified dsDNA templates were *in vitro* transcribed in the presence of natural NTPs to synthesize poly(A) tail-containing sgRNA **1** and **2**, targeting the telomeric repeat and eGFP gene, respectively (Table 1 and Table S1). In order to check if the envisioned sgRNA design was compatible for terminal uridylation, sgRNA **1** was reacted with AMUTP or UTP in the presence of SpCID1, and products were resolved by denaturing PAGE followed by RNA staining (Figure 4). sgRNA **1** showed a slightly diffused band, which can be attributed to transcripts produced with small variations in the number of adenylate residues added at the 3’ end due to known transcript slippage. Rewardingly, bands of lower mobility indicated that the overhang efficiently assisted the terminal uridylation in the presence of natural (**1_UMP_**) and modified (**1_Az_**) UTPs. Similar to uridylation with the model RNA, AMUTP incorporation was found to be biphasic, prominently favoring distributive addition. For downstream applications, the azide-labeled sgRNAs (**1_Az_** and **2_Az_**) were synthesized in large-scale by terminal uridylation of sgRNA **1** and **2** and isolated in good yields (Table 1 and Table S3).

**Figure 4.**
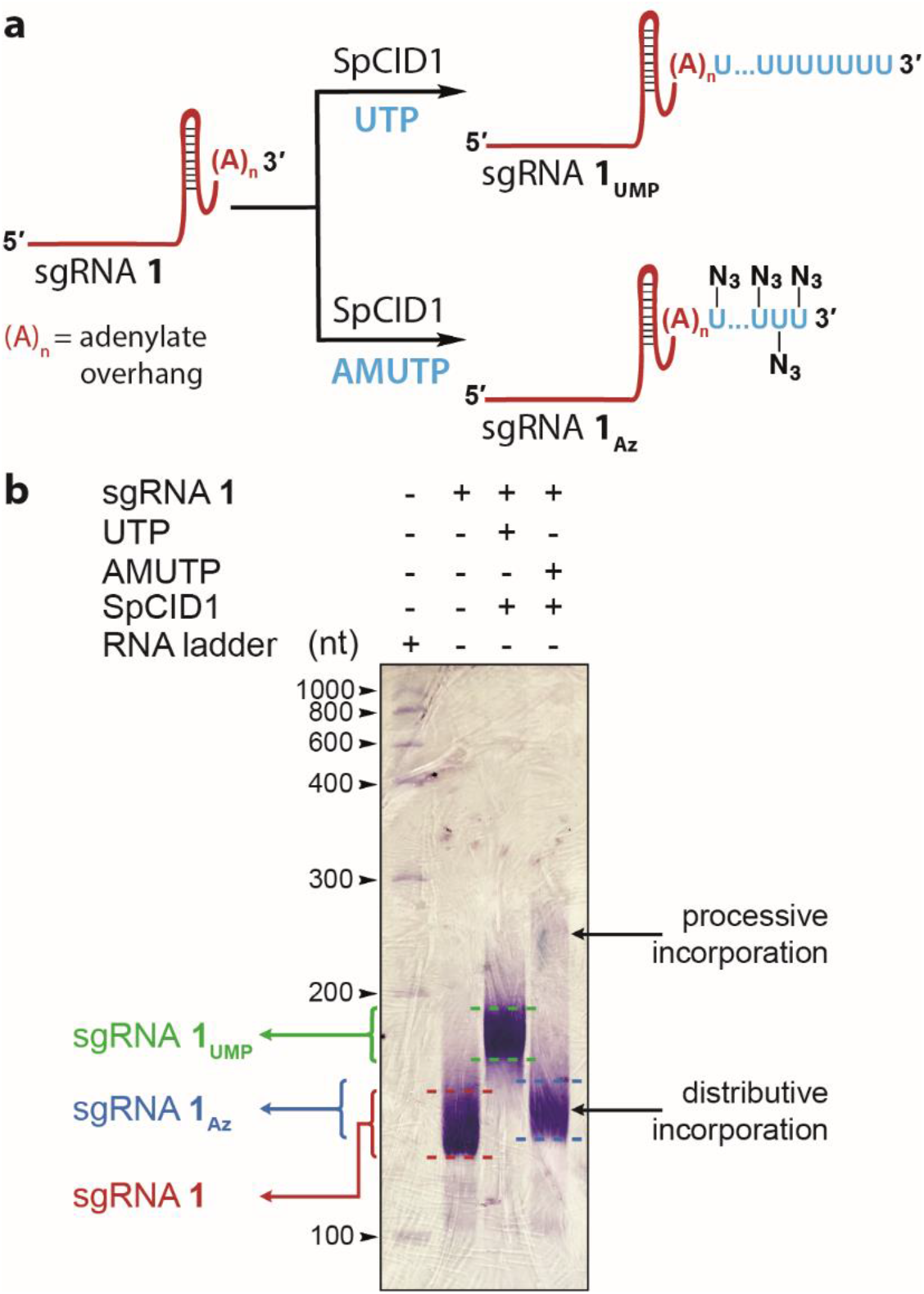
SpCID1 efficiently uridylates poly(A) tail-containing sgRNA in the presence of UTP and AMUTP. **(a)** Scheme showing uridylation reaction setup. **(b)** Uridylated sgRNA products were resolved by PAGE under denaturing conditions and were stained using Stains-All reagent.

### Azide-labeled sgRNAs are compatible for click functionalization

Pre-hybridization click-conjugation of fluorescent and pull-down tags onto sgRNAs was achieved by reacting **1_Az_** and **2_Az_** with strained alkynes Cy3-DBCO and biotin-DBCO (Figure 5a and Figure S7). The reaction products were analyzed by imaging the gel at Cy3 wavelength and or using Stains-All reagent. The staining step revealed the click functionalization of sgRNAs wherein the product bands migrated slowly compared to the azide-labeled sgRNAs produced by distributive and processive incorporation of AMUTP (Figure 5b, Figure S9 and Table S3). In addition, fluorescence image of the gel confirmed the Cy3 labeling of RNAs.

**Figure 5.**
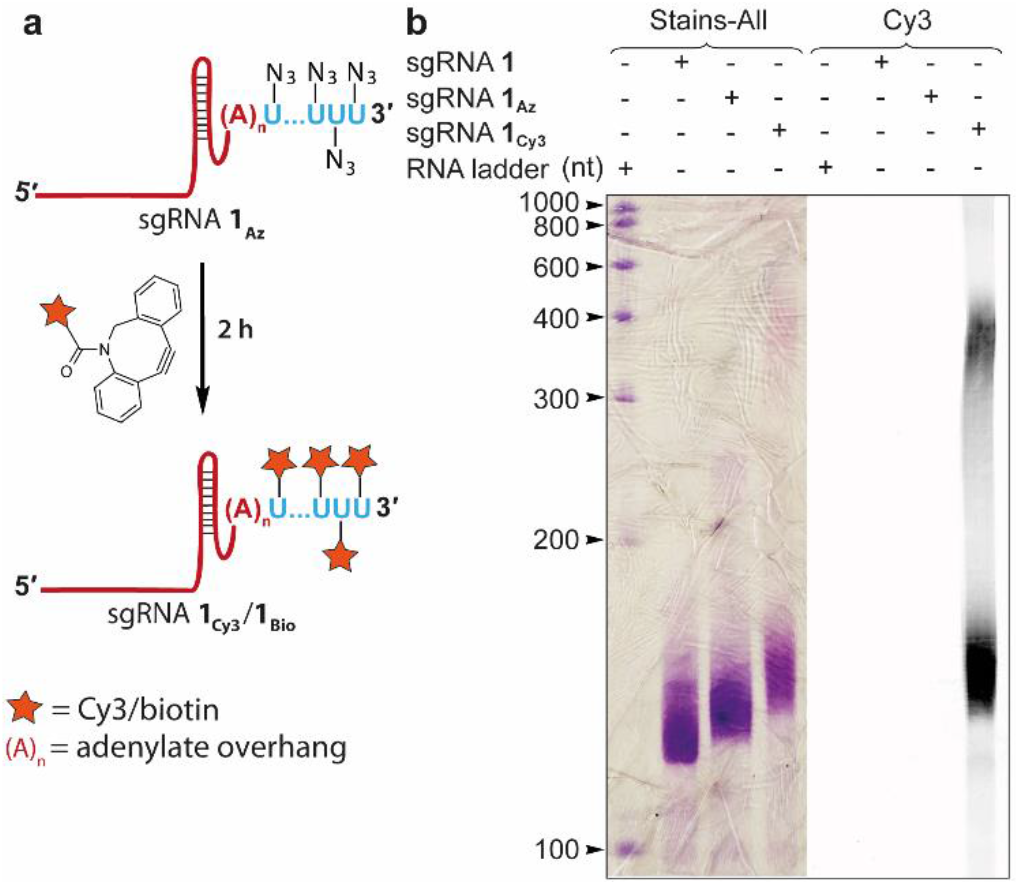
**(a)** Scheme showing click reaction setup. **(b)** Substrate sgRNA and clicked sgRNA product were resolved by PAGE under denaturing conditions. The gel was imaged using a fluorescence scanner (Cy3 wavelength) and then stained using Stains-All reagent. Structure of Cy3-DBCO and biotin-DBCO substrates are shown in Figure S7.

### Modified sgRNAs are functional

To evaluate whether the modified sgRNAs are functional, we performed cleavage reaction using nuclease protein, Cas9 on a target dsDNA (216 bp) corresponding to a segment of eGFP gene. 1:1 molar equivalence of Cas9 and unlabeled sgRNA **2**, azide-labeled (**2_Az_**) or Cy3-conjugated sgRNA (**2_Cy3_**) were incubated at room temperature to form the ribonucleoprotein (RNP) complex. The target dsDNA was added to the complex in the binding buffer, incubated for 30 min at 37 °C and resolved by agarose gel (Figure 6a). The sgRNAs cleaved the dsDNA to produce two fragments of very similar length expected for this target (Figure 6b). sgRNA **2** produced almost complete cleavage under the reaction conditions. While labeled sgRNAs cleaved the target, we observed a progressive reduction in cleavage efficiency in the presence of azide-labeled and click functionalized sgRNAs, which is possibly due to differences in binding affinity of modified sgRNAs to Cas9 or RNP complex to target DNA sequence.

**Figure 6.**
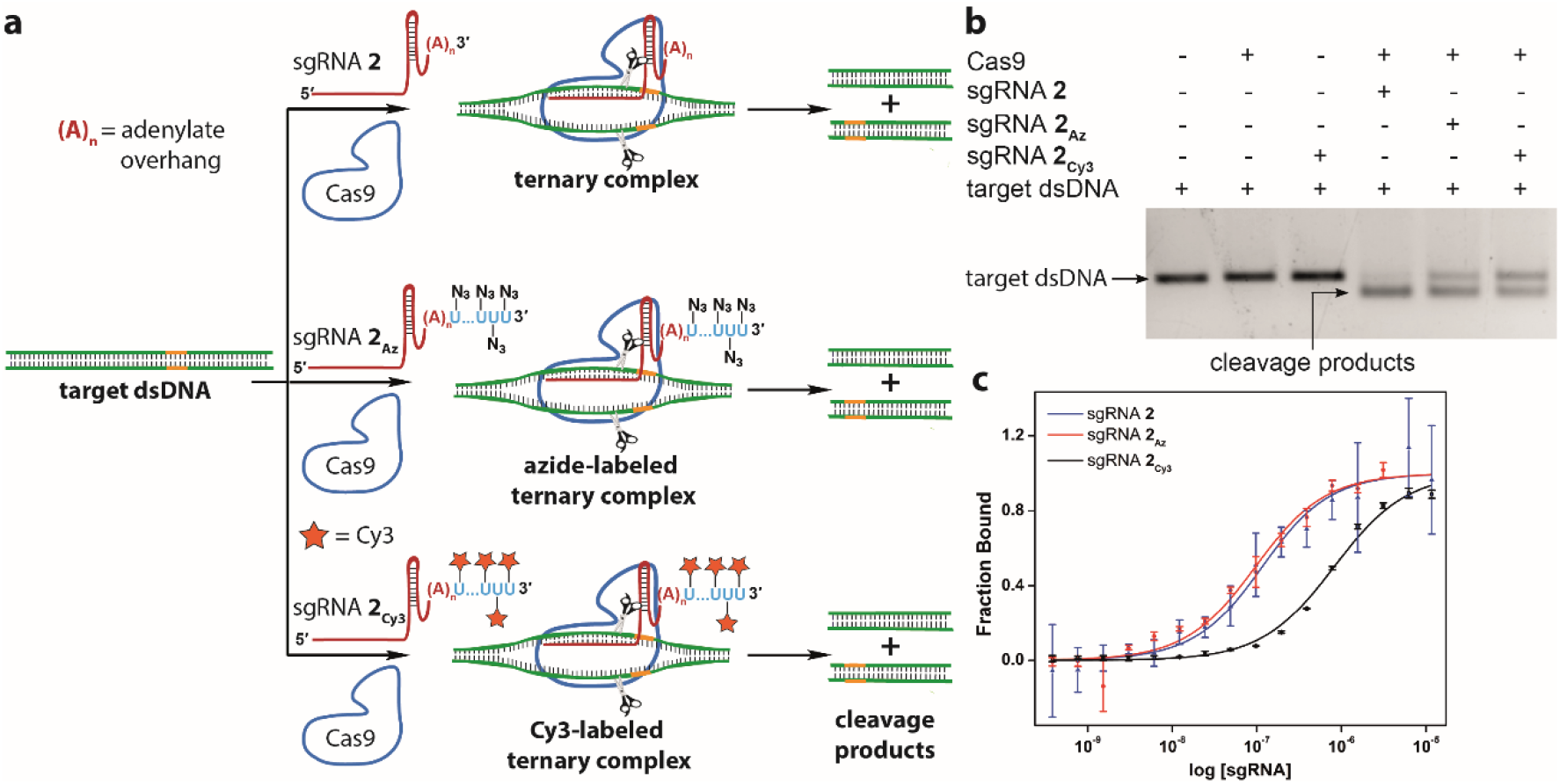
Modified sgRNA-Cas9 complexes cleave target dsDNA. **(a)** eGFP target dsDNA was incubated with Cas9 nuclease protein and sgRNA **2**, azide-labeled sgRNA **2_Az_** or Cy3 click-conjugated sgRNA **2_Cy3_**. **(b)** The cleavage reaction was analyzed by agarose gel electrophoresis, and the bands were stained using EtBr. The target dsDNA in the presence of RNP complex produced two fragments of very similar length consistent with the site of cleavage. **(c)** Curve fits for the binding of unlabeled-**(2)**, AMU-labeled (**2_Az_**) and Cy3 click-labeled (**2_Cy3_**) sgRNAs to dCas9-eGFP derived from MST. Values in c are mean ± s.d for n = 2 independent experiments.

In CRISPR editing or targeting process, sgRNA initially binds to Cas9/dCas9 to form an RNP complex and subsequently this binary complex binds to the target dsDNA^50^. The major structural changes in protein complex occur during the formation of sgRNA-Cas9/sgRNA-dCas9 RNP as compared to RNP-target dsDNA complex^51^. Therefore, any inhibition of RNP complex formation would prevent binding to the target dsDNA and its function. First we determined the efficiency of formation of RNP complex by microscale thermophoresis (MST). dCas9-eGFP fusion protein was titrated with increasing concentration of sgRNA **2**, azide-labeled sgRNA **2_Az_** or Cy3-labeled sgRNA **2_Cy3_** and the binding was monitored by MST using the eGFP tag. A similar apparent dissociation constant (*K_d_*) of 98.1 ± 22.1 nM and 87.4 ± 19.8 nM was observed for the sgRNA **2** and sgRNA **2_Az_** (Figure 6c). The Cy3-labeled sgRNA **2_Cy3_** showed significantly decreased binding affinity (*K_d_* = 799.5 ± 87.6 nM) as compared to the unlabeled and azide-labeled sgRNAs. These results indicate that sgRNA containing minimally perturbing azide tail does not hamper the binary complex formation as opposed to Cy3-clicked sgRNA.

Next, we studied the formation of ternary complex by electrophoretic mobility shift assay (EMSA), which would also help us in determining whether post-hybridization or pre-hybridization click labeling approach is suited for displaying functional tags on target gene, in this case telomeres. sgRNAs (unlabeled **1**, UMP-tailed **1_UMP_**, azide-labeled **1_Az_**, biotin-conjugated **1_Bio_** or Cy3-conjugated **1_Cy3_**) were incubated with dCas9 for 10 min and then with the target dsDNA in a binding buffer (Figure 7a). The dsDNA contained a short telomeric repeat sequence, and one of the strands was labeled with a FAM probe. The samples were resolved by native PAGE and imaged using a gel scanner at FAM and Cy3 wavelengths (Figure 7b and Figure S10). The unlabeled sgRNA **1** showed complete binding to form the ternary complex evident from the formation of a band displaying decreased migration as compared to the dsDNA. On the contrary, UMP-tailed sgRNA showed poor binding. The azide-labeled sgRNA **1_Az_** showed excellent binding of ~85% with very less amount of the unbound fraction. It is to be noted that since AMUTP is incorporated with less efficiency compared to natural UTP in terminal uridylation reaction, there is lesser number of AMUs at the 3’ end of **1_Az_** as compared to Us in **1_UMP_** (*vide supra*, Figure 4b). Hence, UMP-tail on sgRNA could have hindered its binding to dCas9. Therefore, we presume that the lesser incorporation efficiency of AMUTP has in a way assisted the efficient binding of **1_Az_** to dCas9. However, clicked sgRNAs (**1_Bio_** and **1_Cy3_**) showed poor binding efficiency (20–25%) to form the ternary complex potentially due to the introduction of bulky tags (Figure 7b). The formation of small percentage of Cy3-conjugated ternary complex was also evident upon imaging the gel at Cy3 wavelength (Figure S10). Collectively, minimally invasive nature of the azide label complemented by these results indicate that post-hybridization click functionalization approach, i.e., functionalizing ternary complex, would be the method of choice for chemical display of functional tags on gene of interest.

**Figure 7.**
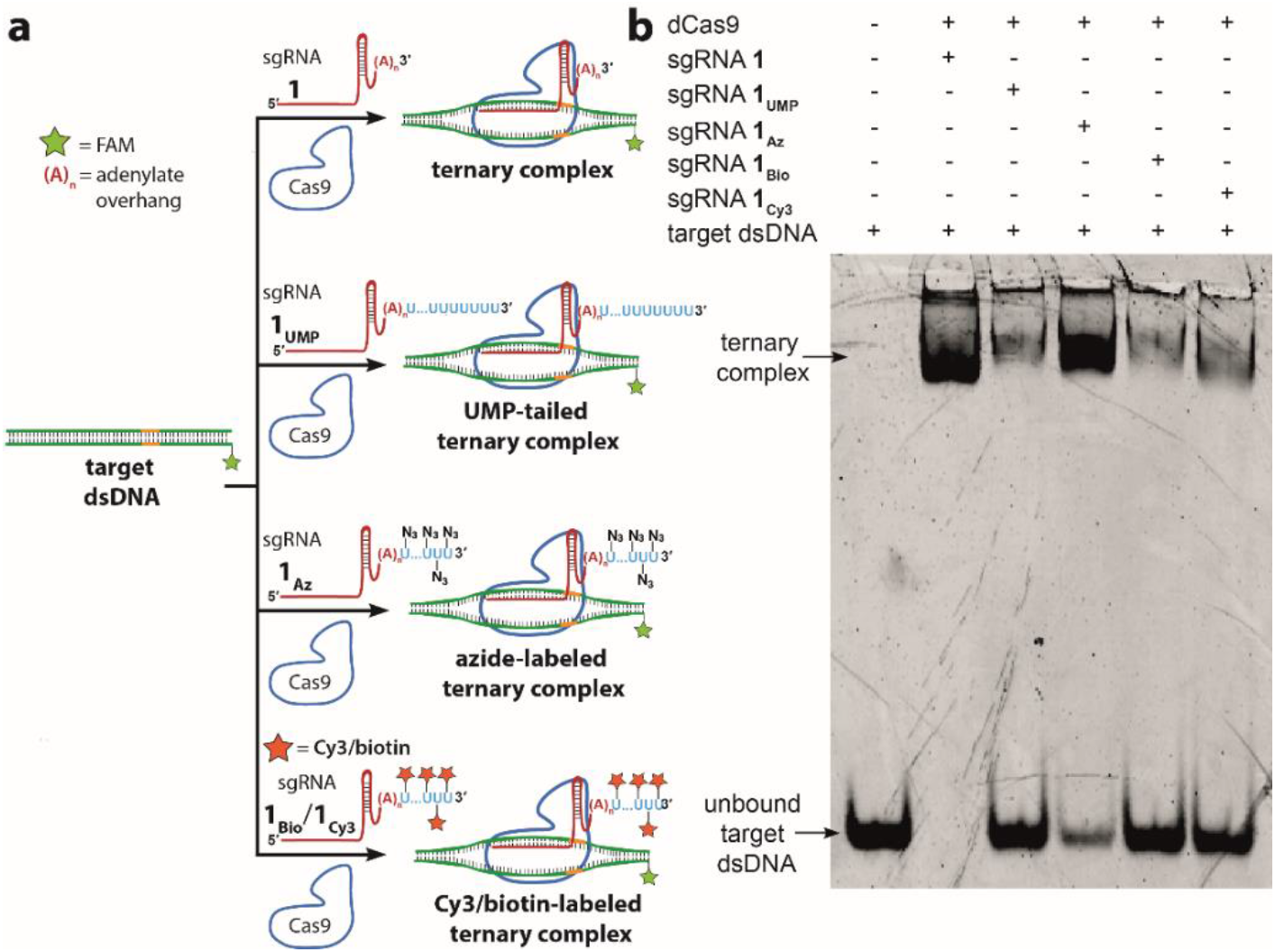
EMSA results suggest that post-hybridization click functionalization is the most favorable approach to display functional tags on target gene. **(a)** Telomeric repeat-containing dsDNA target was incubated with dCas9 and sgRNA **1**, UMP-tailed sgRNA **1_UMP_**, azide-labeled sgRNA **1_Az_**, biotin-conjugated sgRNA **1_Bio_** or Cy3-conjugated sgRNA **1_Cy3_**. The formation of ternary complex (dCas9-sgRNA-dsDNA) was studied by PAGE under native conditions. **(b)** Fluorescence image of the gel at FAM wavelength.

### Azide-tailed sgRNA displays the clickable tags on target gene locus

The localization of azide-labeled sgRNA on the telomeric region of mouse embryonic stem cells (mESCs) was confirmed by CRISPR-FISH^24^. Fixed and permeabilized mESCs were incubated with an RNP complex composed of either unlabeled sgRNA **1’**, azide-labeled sgRNA **1’_Az_** (targeting telomeric region) or azide-labeled sgRNA **2_Az_** (targeting eGFP) and dCas9-eGFP fusion protein (Table 1). sgRNAs, **1’** and **1’_Az_**, were designed to match a validated protospacer sequence used to target telomeric repeat in cells^52,53^. Nuclear puncta corresponding to telomeres were visualized for both **1’** and **1’_Az_**, which is consistent with the telomere localization pattern established using fluorescent dCas9 fusion proteins (Figure 8a, see Figure 8b for single cell image)^12,24,53^. In a control experiment, non-targeting sgRNA **2_Az_** showed fluorescence signal, which was non-specifically distributed all throughout the nucleus. Further, incubation with dCas9-eGFP alone resulted in aggregates mostly localized outside the nucleus. Collectively, these observations confirmed that RNP complex made of azide-tailed sgRNA can be efficiently used to localize clickable groups on the target gene.

**Figure 8.**
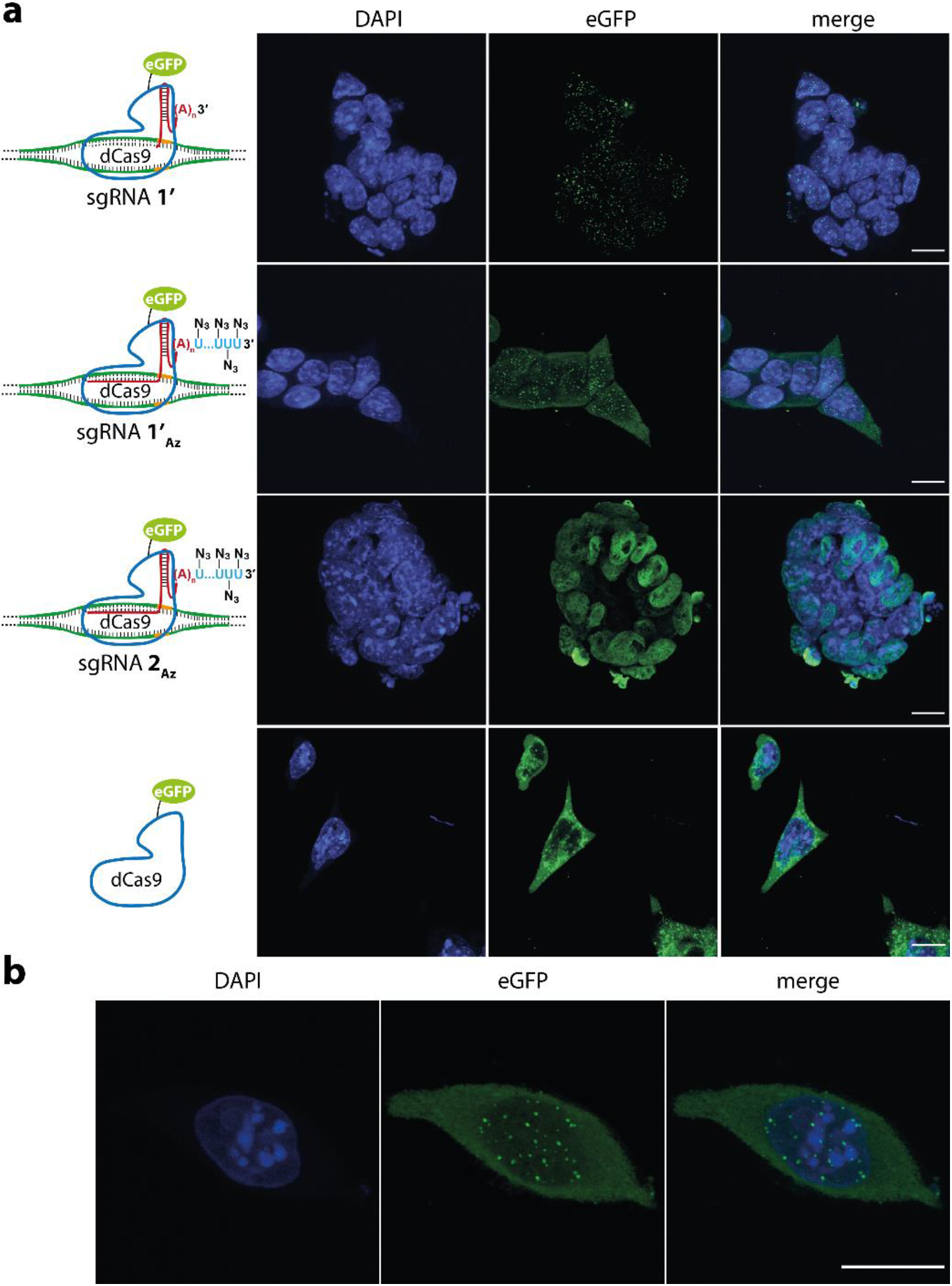
Azide-tailed sgRNA 1’_Az_ localizes on telomeric region. **(a)** mESCs were treated with dCas9-eGFP alone or its RNP complex with unlabeled sgRNA **1’** (targeting telomere), azide-labeled sgRNA **1’_Az_** (targeting telomere) and azide-labeled sgRNA **2_Az_** (targeting eGFP: control). The cells were imaged in DAPI and eGFP channels and maximally projected Z-stacks are shown. First two rows: RNP complex of **1’** and **1’_Az_** directs the localization to telomeric regions visualized as nuclear puncta. Third row: RNP complex of **2_Az_** and dCas9-eGFP non-specifically distributes throughout the nucleus with no observable puncta. Last row: incubation with dCas9-eGFP alone resulted in aggregates, which were mostly localized outside the nucleus (scale bar, 10 μm). **(b)** Maximally projected Z-stack image showing nuclear puncta localized to telomeres in a single embryonic stem cell obtained using RNP complex of **1’_Az_** and dCas9-eGFP (scale bar, 10 μm). The experiments were performed as n = 2 biological replicates.

### sgR-CLK enables site-directed functional tagging of target gene

To evaluate the compatibility of azide tags localized on the telomeric region for click-functionalization we performed CuAAC and SPAAC reactions on the isolated chromatin to enrich the target gene (Figure 9a). Fixed and permeabilized mESCs were incubated with azide-labeled sgRNA **1’_Az_** and dCas9-eGFP complex. The localization of azide labels on the telomere region was initially confirmed by the presence of nuclear puncta in the eGFP channel (Figure 9a). The cells were then cross-linked and the nuclear isolate was sheared until chromatin fragments of the range of 200–500 bp were obtained. CuAAC reaction was performed on chromatin using a biotin-alkyne in the presence of a catalytic system formed by CuSO_4_, sodium ascorbate and THPTA. In case of SPAAC reaction, chromatin was first treated with iodoacetamide to sequester protein thiols on chromatin so as to reduce non-specific enrichment resulting from the reaction between the strained biotin-alkyne substrate and thiols^54^. Further, strained alkyne system, sDIBO was preferred in this reaction as DBCO-based alkynes are known to show non-specific reaction with cellular thiols (Figure S7)^54^. After click reaction, unreacted biotin substrate was removed and biotinylated-chromatin was incubated with streptavidin-coated magnetic beads. The captured chromatin was eluted, de-crosslinked and the isolated genomic DNA fragments were subjected to qPCR using telomeric primers as previously described^55^. In parallel, a control azide-labeled sgRNA **2_Az_** was subjected to same experimental procedure to account for the background enrichment from non-target sgRNA and click reaction. sgR-CLK in the presence of **1’_Az_** revealed significant enrichment of the telomere region by both CuAAC and SPAAC reactions (Figure 9b). On the other hand, chromatin captured using nontargeting sgRNA **2_Az_** by click reactions showed very low enrichment. Further, fold enrichment of telomeric repeat region calculated for sgRNA **1’_Az_** upon normalizing with sgRNA **2_Az_** was found to be nearly 12 and 6 while employing CuAAC and SPAAC reactions (Figure 9c). Notably, CuAAC reaction exhibited better pull-down efficiency as compared to SPAAC reaction, possibly due to better accessibility and reactivity of the terminal alkyne substrate as compared to the bulky cyclooctyne counterpart^56^. It is important to mention here that sgR-CLK enriched telomere region discernibly higher (12-fold) as compared to telomere capture using fusion proteins and MS2 aptamer (~5-fold)^11,26^. Taken together, these results indicate that sgR-CLK is a robust technique for site-directed functionalization of chromatin, and is expected to expand the utility of CRISPR-dCas9 as a gene targeting tool.

**Figure 9.**
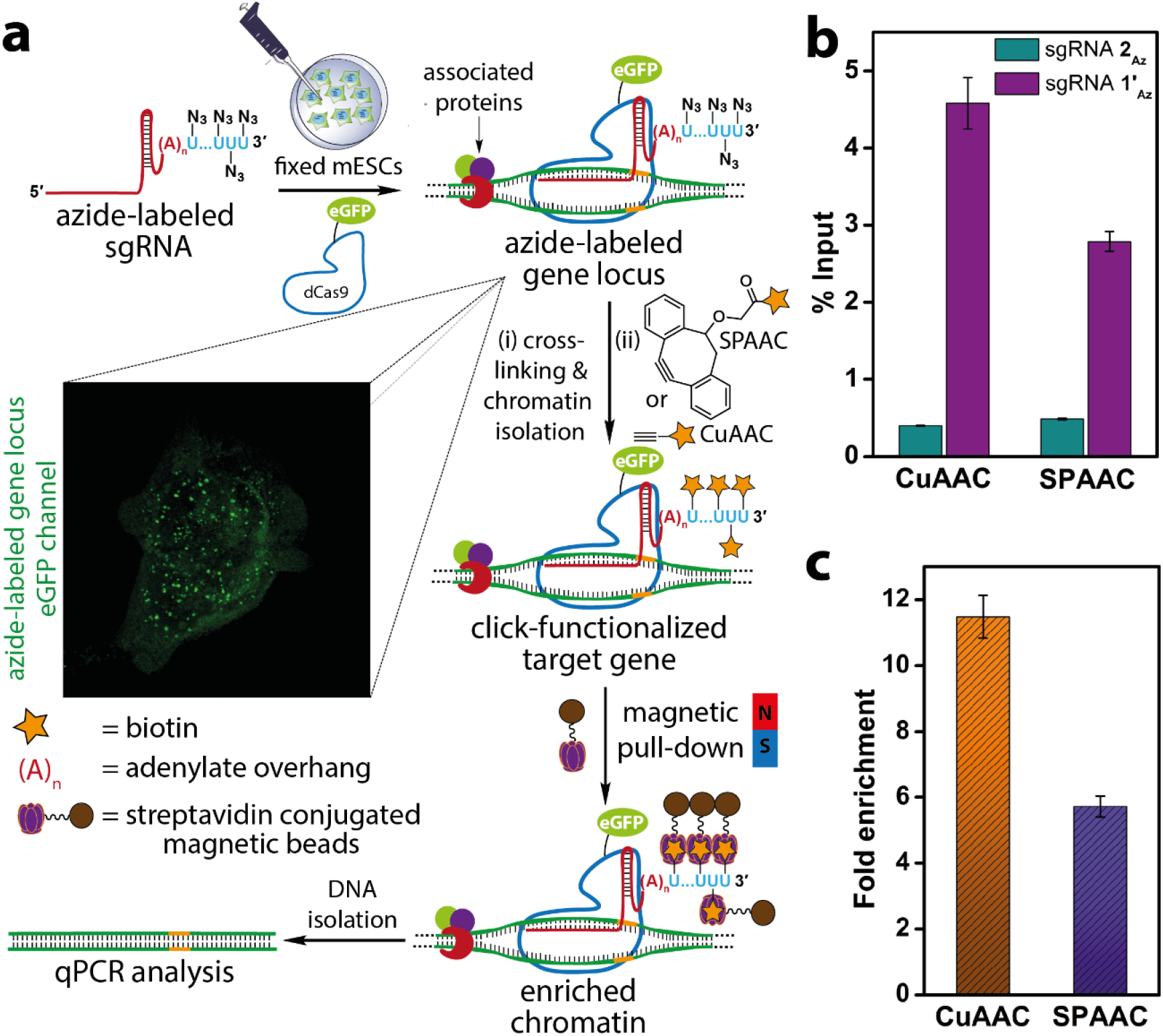
Site-directed functionalization of telomeres by sgR-CLK was confirmed by ChIP-qPCR. **(a)** Schematic diagram illustrating the steps involved in chromatin capture by *in situ* click reaction performed on the target-bound ternary complex using biotin-alkynes. While CuAAC reaction was performed using a biotin substrate containing a terminal alkyne, SPAAC reaction was performed using biotin-conjugated to a strained alkyne (sDIBO). See Figure S7 for complete structure of alkyne substrates. Biotinylated chromatin was enriched using streptavidin beads and subjected to qPCR analysis. **(b)** qPCR analysis of enriched chromatin obtained by CuAAC and SPAAC reactions using telomere-targeting sgRNA **1’_Az_** and control non-targeting sgRNA **2_Az_**. The enrichment is expressed relative to respective inputs after click reaction step. **(c)** A plot showing the fold enrichment of telomere DNA using sgRNA **1’_AZ_** normalized over sgRNA **2_Az_** for CuAAC and SPAAC reactions (see Methods section for details). Values in b and c are mean ± s.d. performed as n = 3 technical replicates.

## Discussion

Several CRISPR-dCas9-based methods have recently emerged as powerful tools for localizing genetically encodable biomolecules to manipulate target genomes. Typically, these methods use expression platforms to recruit payloads on dCas9, which are then directed by sgRNA to the target gene via the formation of a ternary complex^18^. Alternatively, sgRNA scaffolds containing sequences for binding proteins have offered systems to activate or repress gene and also visualize specific gene loci^17,19,57^. However, analogous tools for gene-specific display of chemical cargos are much less developed due to paucity of efficient labeling technologies. Therefore, development of a modular technology that would enable gene-specific localization of synthetic recruits of choice would immensely expand the repertoire of CRISPR as a gene editing and targeting tool. To this end, we developed a technique called sgR-CLK to recruit functional tags on the target gene by combining the power of bioorthogonal azide-alkyne click chemistry and our ability to engineer sgRNAs using a uridylyl transferase enzyme (SpCID1).

Based on the structure of SpCID1 bound to the nucleotide and its ability to polyuridylate unstructured poly(A) tail containing RNA, we configured sgRNAs to contain a short poly(A) unit at the 3’ end. Rewardingly, this design enabled us to repurpose SpCID1 to introduce clickable azide tags on sgRNAs in the presence of an azide-modified UTP analog, namely AMUTP. The azide-tailed and corresponding click functionalized sgRNAs complexed with Cas9 were found to be functional as the RNP complexes cleaved the target dsDNA with reasonable efficiency. Importantly, we noted that the azide-tailed sgRNA formed CRISPR ternary complex almost as efficiently as the unlabeled sgRNA. However, biotin- and Cy3-clicked sgRNA constructs formed very less of the ternary complex, alluding that clicking ternary complex containing minimally invasive azide handles is a better strategy to recruit functions on the target gene. Indeed, an azide-tailed sgRNA targeting the telomeric DNA repeat exhibited nuclear puncta corresponding to the telomere region in embryonic stem cells, which indicated the locus-specific display of azide groups on the target gene. Further, click reactions performed on chromatin using biotin alkynes, followed by chromatic capture revealed significant fold enrichment of telomeres, which depended on whether copper-catalyzed or strain-promoted reaction was employed. Notably, sgR-CLK under CuAAC reaction conditions produced better enrichment of the target gene in comparison to other CRISPR-based telomere capture methods using fusion proteins and MS2 aptamer. These results clearly validate the key concept of this study, i.e., directing clickable groups to a specific region of chromatin, which can further enable the display of synthetic recruits by sgR-CLK.

This technology has several advantages. First, it is modular on two counts. Our design enables insertion of multiple azide groups at the 3’ end of any sgRNA sequence. In addition, TUTase can be used to incorporate other minimally perturbing bioorthogonal reactive groups such as alkyne, vinyl and cyclopropene, which would allow introduction of a range of functions by using cognate reaction partner^58^. Hence, a combination of these two features can be conceivably used to prepare clickable sgRNAs targeting same or different gene loci to either achieve a robust functional output at a designated genomic site or interrogate multiple genes in parallel^59^. Another layer of foreseeable utility is the mutimeric display of small molecules (e.g., drugs or probes) on the target gene, which could enhance the efficacy and specific action of the synthetic cargo. Although in this work, sgR-CLK has been demonstrated as a proof-of-concept in fixed cells, we would be extending this technology to live cells for multimeric locus-specific display of synthetic molecules including epigenetic modulators and transcriptional activator/repressors, which are otherwise accomplished by less versatile approaches using DNA binding proteins or pyrrole-imidazole polyamides^29,60,61^. On the whole, sgR-CLK is anticipated to open up new experimental strategies for gene-specific display of chemical cargos.

## Methods

Detailed procedure for cloning, expressing SpCID1, single nucleotide incorporation, click labeling of RNA is described in Supplementary Information.

### Uridylation of RNA oligonucleotide using SpCID1

Model 5’-FAM-labeled RNA oligonucleotide (10 μM, Table 1) was incubated with UTP, AMUTP, APUTP or ATUTP (500 μM) in the presence of Tris-HCl buffer (10 mM, pH 7.9 at 25 °C), NaCl (50 mM), MgCl_2_ (10 mM), DTT (2 mM), RiboLock RNase inhibitor (1 U/μL) and 1 μL of SpCID1 (10.25 pmol) in a final volume of 20 μL. After 5, 15 and 30 min, 5 μL aliquots of reaction mixture (50 pmol of RNA oligonucleotide) were mixed with 15 μL of denaturing loading buffer (7 M urea in 10 mM Tris-HCl, 100 mM EDTA, 0.05% bromophenol blue, pH 8) and heat-denatured at 75 °C for 3 min. 5 μL of the sample was loaded on to 20% denaturing polyacrylamide gel, electrophoresed and imaged using Typhoon gel scanner at FAM wavelength.

### Constructing sgRNAs containing 3’-adenylate overhang

Forward (CFP1, CFP1’ or CFP2) and reverse CRP primers (1 μM) were incubated with sgRNA templates CT1, CT1’ or CT2 (44 nM) in EmeraldAMP GT PCR master mix in a final volume of 25 μL. PCR conditions: heat denaturation at 94 °C for 1 min, 35 cycles of (denaturing: 94 °C for 20 s, annealing: 64 °C for 20 s, extension: 68 °C for 30 s), final extension at 68 °C for 5 min. Multiple small-scale PCRs were performed to isolate dsDNA templates required for large-scale transcription reaction. The amplicons were purified using NucleoSpin^®^ Gel and PCR Clean-up kit. PCR with CT1, CT1’ and CT2 yielded dsDNA templates for the synthesis of sgRNAs **1, 1’** and **2**. For primer and template sequences see Table S1.

*In vitro* transcription reactions were performed using ATP, GTP, CTP and UTP (2 mM), RiboLock RNase inhibitor (0.4 U/μL), respective dsDNA templates (300 nM) and T7 RNA polymerase (3.2 U/μL) in 40 mM Tris-HCl (pH 7.9), 10 mM MgCl_2_ 20 mM DTT, 10 mM NaCl and 1 mM spermidine in a final reaction volume of 250 μL. The reaction was incubated at 37 °C for 12 h. 3’-adenylate sgRNAs **1, 1’** and **2** were isolated by MEGAclear™ Transcription Clean-Up kit using manufacture’s protocol. A typical 250 μL transcription reaction yielded 2–3 nmol of the sgRNAs.

### Azide-tailed sgRNAs using SpCID1

*In vitro* transcribed sgRNA **1** (10 μM, 200 pmol) was incubated with natural UTP or AMUTP (500 μM) in the presence of Tris-HCl buffer (10 mM, pH 7.9 at 25 °C), NaCl (50 mM), MgCl_2_ (10 mM), DTT (2 mM), RiboLock RNase inhibitor (1 U/μL) and SpCID1 (20.5 pmol) in a final volume of 20 μL. The reaction was incubated at 37 °C for 30 min and further heat-denatured by incubating at 75 °C for 3 min. The reaction mixture (150 pmol) was resolved in 8.5% denaturing polyacrylamide gel. Bands corresponding to UMP-tailed sgRNA **1_UMP_** and azide-labeled sgRNA **1_Az_** were visualized by Stains-All reagent (Figure 4).

Multiple such small-scale (20 μL, 200 pmol of sgRNA) reactions were performed with **1, 1’** and **2** and the reactions were pooled together and the product sgRNAs **1_Az_, 1’_Az_** and **2_Az_** were precipitated by addition of 1 volume of 5 M ammonium acetate and 10 volume of ethanol followed by incubation for 2 h at −20 °C. The samples were centrifuged at 15,000 rpm for 15 min and RNA pellets obtained were washed with chilled 75% ethanol in water and sgRNAs **1_Az_, 1’_Az_** and **2_Az_** thus obtained were dissolved in autoclaved water. See Table S3 for yields.

### *In vitro* cleavage assay

A 216 bp dsDNA template corresponding to a segment of eGFP gene required for cleavage assay was generated by PCR using forward (eGFPF) and reverse (eGFPR) primers (0.5 μM). The primers in the presence of dNTPs (0.2 mM) were incubated with linearized eGFP-N1 plasmid (100 ng) and Taq DNA polymerase (2.5 U) in 1X PCR buffer in a final volume of 50 μL. PCR conditions: heat denaturation at 95 °C for 5 min, 35 cycles of (denaturing: 95 °C for 30 s, annealing: 58 °C for 30 s, extension: 72 °C for 30 s) and final extension at 72 °C for 5 min. The amplicons were purified by using PCR Clean-up kit.

A 1:1 solution (1.25 μM) of sgRNA **2**, azide-labeled sgRNA **2_Az_** or Cy3-labeled sgRNA **2_Cy3_** and Cas9 was incubated at room temperature for 10 min. eGFP dsDNA was mixed with the above RNP complexes in 20 mM HEPES (pH 7.5), 100 mM KCl, 5 mM MgCl_2_, 1 mM DTT to make a final volume of 15 μL. The final concentration of dsDNA and RNP complex was 50 nM and 250 nM, respectively. The reaction was performed for 30 min at 37 °C. Proteinase K (40 μg) was added to the reaction mix and incubated at 55 °C for 30 min. Proteinase K was deactivated by incubating at 70 °C for 10 min and RNase A (10 μg) was added. DNA gel loading buffer was added and the samples were loaded on a 2% agarose gel, resolved and stained using EtBr (Figure 6b).

### MST analysis to quantify RNP complex^62^

dCas9-eGFP (180 nM) in 20 mM HEPES (pH 7.5), 150 mM KCl, 10 mM MgCl_2_ and 1 mM DTT was incubated with varying concentrations of sgRNA **2**, azide-labeled sgRNA **2_Az_** or Cy3-labeled sgRNA **2_Cy3_** (0.4 nM to 12.5 μM) in a final volume of 10 μL for 10 min at room temperature. The samples (~5 μL) were loaded onto standard treated quartz capillary and excited at 60% LED power and 40% MST power at 25 °C using blue filter, and MST was recorded. The experiments were performed individually as two replicates. Data was analyzed using MST analysis software to determine bound protein fraction with respect to sgRNA concentration.

### EMSA to estimate ternary complex formation

sgRNA **1, 1_UMP_, 1_Az_, 1_Bio_** or **1_Cy3_** (2 μM) was incubated with dCas9 (2.5 μM) in the presence of RiboLock RNase inhibitor (4 U/μL) in a total volume of 5 μL at room temperature for 10 min. 5 μL solution of target dsDNA (0.2 μM, formed by annealing TS1 and TS2 DNA oligonucleotides, Table S1) and DTT (20 μM) was added to the individual RNP complex in binding buffer (20 mM HEPES (pH 7.5), 100 mM KCl, 5 mM MgCl_2_, 1 mM DTT, 1 mg/mL BSA, 0.1% Triton X-100, 5% glycerol). The samples were incubated for 30 min at 37 °C. The samples (0.5 pmol DNA) were resolved by native PAGE (8% gel containing 10 mM MgCl_2_) at 4 °C in 1X TBE buffer supplemented with 10 mM MgCl_2_. The gel was imaged using Typhoon gel scanner at FAM wavelength (Figure 7).

### Cellular localization of azide-labeled sgRNA on telomeres

mESCs (R1/E) were seeded in coverslips and cultured in DMEM in the presence of leukemia inhibiting factor (LIF) at 37 °C and 5% CO_2_ to maintain cells in its pluripotent state^63^. Post-seeding (24 h), the media was removed and cells were washed with 1X PBS. The cells were then fixed in pre-chilled 50% acetic acid in methanol at −20 °C for 20 min. The cells were washed three times with 1X PBS for 5 min with gentle shaking. Further, the cells were permeabilized by incubating in hybridization buffer (20 mM HEPES (pH 7.5), 150 mM KCl, 10 mM MgCl_2_ 1 mM DTT, 2% BSA, 5% glycerol and 0.1% Triton-X) at 37 °C for 30 min. dCas9-eGFP (200 nM) and sgRNA **1**, sgRNA **1’_Az_** or sgRNA **2_Az_** (200 nM) and RiboLock RNase inhibitor (1.6 U/μL), were incubated for 20 min at room temperature in hybridization buffer in a final volume of 50 μL. The cells were incubated with the respective RNP complex or dCas9-eGFP alone in a humid chamber for 4 h at 37 °C and then washed three times with hybridization buffer for 5 min. The cells were stained with 5 μg/mL DAPI and the coverslip was mounted on glass slides for imaging. The cells were visualized in Leica TCS SP8 confocal microscope in DAPI and eGFP channels. Images were deconvoluted on Leica LAS X software using blind deconvolution and Z-stacks were projected (maximum intensity) using Fiji image analysis software (Figure 8).

### Chromatin capture using sgR-CLK

mESCs (R1/E) were seeded in 10 cm dish and cultured in DMEM in presence of leukemia inhibiting factor (LIF) at 37 °C and 5% CO_2_. At around 70% confluency, cells were fixed and permeabilized as above. dCas9-eGFP (200 nM), azide-labeled sgRNA **1’_Az_** or **2_Az_** (200 nM) and SUPERaseIn RNAse Inhibitor (0.1 U/μL) were incubated in hybridization buffer for 20 min at room temperature in a final volume of 1 mL for the formation of RNP complex. The cells were incubated with the above solution in a humid chamber for 1.5 h at 37 °C and then washed with 1X PBS for 5 min. Cells were cross-linked with 1% formaldehyde in DMEM for 10 min at room temperature with gentle shaking. Further, glycine (125 mM) was added and incubated for 5 min. Cells were harvested and washed twice with 1X protease inhibitor cocktail (PIC) in 1X PBS. For nuclear isolation, cells were resuspended in 2 ml of PBS followed by the addition of 2 mL of nuclear isolation buffer (40 mM Tris HCl pH 7.5, 1.28 M sucrose, 20 mM MgCl_2_ and 4% Triton X-100) in a final volume of 10 mL. Cell suspension was incubated on ice for 20 min with frequent mixing. The nuclear isolate was centrifuged at 2500 g for 15 min at 4 °C. The nuclear pellet obtained was resuspended in 1 mL RNA immunoprecipitation buffer (25 mM Tris HCl pH 7.5, 150 mM KCl, 5 mM EDTA pH 8.0, 0.5 mM DTT, 0.5% Igpal, 0.1 U/μL SUPERaseIn RNAse Inhibitor, 1X PIC) and sonicated for 4 cycles of 15 min (30 s on and 30 s off) at 4 °C in a Bioruptor^®^ Plus sonication device. After sonication, the fragment size (200-500 bp) was confirmed by agarose gel electrophoresis. Further, the nuclear debris were centrifuged at 13,300 rpm for 10 min and supernatant was transferred to a fresh vial to afford the chromatin. For SPAAC reaction, chromatin was treated with 50 mM iodoacetamide in a final volume of 1 mL for 60 min at room temperature. Samples were buffer-exchanged to 1X PBS using 10 kDa cut-off columns at 4 °C. SPAAC reaction on chromatin was performed with 100 μM biotin-sDIBO alkyne in a final volume of 500 μL having 0.1 U/μL SUPERaseIn RNAse Inhibitor. CuAAC reaction on chromatin was performed with 100 μM biotin-alkyne in the presence of 1 mM CuSO_4_, 1 mM sodium ascorbate, 1 mM THPTA. Both the reactions were performed for 1 h at room temperature. The excess biotin was removed by buffer exchanging to 1X PBS using 10 kDa size cut-off columns to a final volume of 1 mL.

Meanwhile, 50 μL of Dynabeads MyOne Streptavidin C1 magnetic beads for each sample of the click reaction was washed thrice with 1X PBS. 5% volume inputs (50 μL) were taken out, stored at −20 °C and remaining samples were treated with 50 μL of Dynabeads, 0.1 U/μL SUPERaseIn RNAse Inhibitor and incubated overnight with gentle rotation at 4 °C. The immobilized samples were washed with 1 mL of each wash buffer for 5 min at 4 °C with gentle rotation (once with low salt buffer: 20 mM Tris HCl (pH 8.0), 150 mM NaCl, 2 mM EDTA, 1% Triton X-100, 0.1% SDS; once with high salt buffer: 20 mM Tris HCl (pH 8.0), 500 mM NaCl, 2 mM EDTA, 1% Triton X-100, 0.1% SDS; once with LiCl buffer: 10 mM Tris HCl (pH 8.0), 1 mM EDTA (pH 8.0), 250 mM LiCl, 1% sodium deoxycholate, 1% Igpal; twice with TE buffer: 10 mM Tris HCl (pH 8.0), 1 mM EDTA).

The chromatin from each sample was eluted from the beads by incubating with 500 μL of elution buffer (100 NaHCO_3_, 1% SDS) at room temperature for 15 min. Input samples were also treated with 500 μL of elution buffer under same conditions. The eluted chromatin samples were de-crosslinked by incubating in 200 mM NaCl at 65 °C overnight. Each sample was treated with proteinase K (20 μg), 10 μL of 0.5 M EDTA (pH 8.0) and 20 μL of 1 M Tris HCl (pH 7.5) at 42 °C for 60 min for digesting the proteins. Genomic DNA fragments from each sample were isolated using PCR-clean up kit subjected to qPCR.

### qPCR to estimate telomere DNA enrichment

To 2 μL of 1% of input or 2 μL of pull-down samples corresponding to azide-labeled sgRNA **1’_Az_** and **2_Az_** obtained by SPAAC and CuAAC reactions were added 7.5 μL TB Green Premix Ex Taq II and 1 μL TeloF and TeloR primer (0.5 μM each, Table S1). The final volume was adjusted to 15 μL with water. qPCR conditions: heat denaturation at 95 °C for 1 min, 40 cycles of (denaturing: 95 °C for 10 s, annealing: 52 °C for 30 s, extension: 72 °C for 30 s), final extension at 72 °C for 5 min. The Ct values for input were adjusted to 100% and then pull-down values were normalized with 100% input and represented as enrichment relative to respective inputs for sgRNA **1’_Az_** and sgRNA **2_Az_** for CuAAC and SPAAC reactions, respectively. Further, the fold enrichment for telomere-pull down with sgRNA **1’_Az_** over non-targeting sgRNA **2_Az_** was calculated for both the click reactions using the equation given below. Reactions were performed in triplicate.

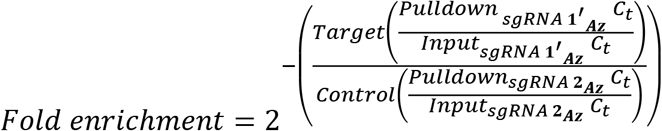

## Acknowledgements

We thank Dr. Gayathri Pananghat and Dr. Amrita Hazra for their help in cloning and protein purification experiments. We thank Rupam Bhattacharjee for his help in the mass analysis of RNA oligonucleotides. J.T.G. thanks IISER Pune and Wellcome Trust-DBT India Alliance for graduate research fellowship. D.C. acknowledges research grant from DBT (GAP 0175). This work was supported by Wellcome Trust-DBT India Alliance (IA/S/16/1/502360) to S.G.S.

## Author contributions

S.G.S. and J.T.G. designed the study. J.T.G. synthesized the nucleotide analogs and performed cloning, protein purification and established sgR-CLK. M. Azhar and J.T.G. performed ChIP and qPCR assays. M. Aich performed FISH experiments. D.S. performed MST analysis. U.A. assisted J.T.G. in performing cloning and protein purification experiments. S.M. and D.C. guided in establishing MST, CAS-FISH, ChIP assays to validate sgR-CLK. All the authors analyzed the data, and J.T.G. and S.G.S. wrote the manuscript in consultation with all the authors.

## Additional Information

Supplementary information is available.

