## Supplementary Information for "TUTase mediated site-directed access to clickable chromatin employing CRISPR-dCas9"

### Materials

*Schizosaccharomyces pombe* was gifted by Dr. Devyani Haldar (CDFD, Hyderabad). Phusion® High-Fidelity DNA Polymerase, Dpn1 was acquired from New England Biolabs. DNA oligonucleotides (ONs) were purchased from Integrated DNA Technologie, Inc. and Sigma Aldrich. NTPs, T7 RNA polymerase, ribonuclease inhibitor (RiboLock), SUPERase In™ RNase Inhibitor, Low Range RNA Ladder, Click-iT™ biotin sDIBO alkyne, Alexa Fluor™ 594 alkyne, Superscript III, TRIzol, Dynabeads™ MyOne™ Streptavidin C1, MEGAclear™ transcription Clean-Up kit, DMEM, SYBR™ Safe Stain, RNA ladder were obtained from Thermo Fischer Scientific. PrimeSTAR GXL DNA Polymerase, EmeraldAMP GT PCR master mix, TB Green™ Premix Ex Taq™ II and DNA ladders were purchased from TakaraBio. DH5α, BL21 (DE3) and Rosetta™(DE3) competent cells were provided by Dr. Thomas J. Pucadyil (IISER Pune). Taq DNA polymerase was purchased from Geneaid. pET28a plasmid was acquired from Novagene. The modified triphosphates used, AMUTP (Commercially made available with JenaBiosciences), APUTP, ATUTP were synthesized as per our reported procedure.[S1] dCas9 was purchased from PNABio whereas Cas9 and dCas9-eGFP were expressed.[S2] Biotin-PEG4-DBCO (biotin-DBCO), Cy3-DBCO and biotin-PEG4-alkyne (biotin-alkyne) were obtained from Click Chemistry Tools. Stains-All, iodoacetamide and reagents for buffer preparation (BioUltra grade) were acquired from

Sigma Aldrich. Luria Broth, agar, yeast extract were purchased from HiMedia. Vivaspin 20 (10 kDa) molecular-weight cut-off columns were acquired from GE Healthcare. NucleoSpin® PCR Clean-up kit and QIAquick PCR purification kit were purchased from Macherey-Nagel and Qiagen. cOmplete EDTA-free protease inhibitor cocktail was purchased from Roche. Standard treated quartz capillary for MST was acquired from nanoTEMPER technologies. All experiments were performed using autoclaved water.

**Instrumentation**

Oligonucleotides and transcripts were quantified by measuring absorbance at 260 nm in UV-2600 Shimadzu or NanoDrop™ 2000c spectrophotometer. Labeled RNA oligonucleotides were purified using Agilent Technologies 1260 Infinity HPLC. ESI-MS analysis of RNA oligonucleotides was performed using Waters SYNAPT G2-Si Mass Spectrometry instrument in negative mode. Bacterial cultures were incubated in New Brunswick Innova 4230 Refrigerated Incubator Shaker and Thermo Scientific Forma Orbital Shaker 480. Size exclusion chromatography was performed on Biorad NGC™ Chromatography system on a HiLoad 16/600 Superdex 200 preparative column. PCR was performed on Mastercycler® pro PCR machine from Eppendorf. Oligonucleotides were resolved by PAGE using an OWL S4S sequencing gel electrophoresis instrument and were imaged using Typhoon TRIO+ Variable mode Imager. Microscale thermophoresis (MST) was measured in Monolith NT.115 from nanoTEMPER Technologies. Sonication for chromatin precipitation was performed in Bioruptor® Plus sonication device from Diagenode. qPCR was performed in LightCycler® 480 Instrument II from Roche.

**Cloning of SpCID1 gene into bacterial expression vector**

Fission yeast, *Schizosaccharomyces pombe* was grown in YES media (100 mL) for 24 h at 28 °C until an $OD_{600}$ of ~1.4 was reached. The cells were pelleted down and the total RNA was extracted using TRIzol and its integrity was confirmed by agarose gel (1.5%) electrophoresis in 1X TAE buffer. Cellular RNA obtained (1.9 µg) was incubated with oligo$(dT)_{20}$ primer (5 µM), dNTPs (1 mM) in water in a final volume of 10 µL (Figure S1a, S2 and Table S1). The cocktail was heated at 65 °C for 5 min and flashed cooled to 4 °C in 30 s for primer annealing. To the reaction mix, $MgCl_2$ (5 mM), DTT (0.01 mM), RNAse OUT (2 U/µL), Superscript III (10 U/µL) and 2 µL of 10X reverse transcription buffer (200 mM Tris-HCl, 500 mM KCl, pH 8.4) was added in a final volume of 20 μL. The solution was incubated at 25 °C for 5 min and cDNA synthesis was performed at 50 °C for 50 min. Further the reverse transcriptase enzyme was heat denatured at 85 °C for 5 min and sample cooled to 4 °C. To the reaction, 1 μL of *E. coli* RNase H (2 U) was added and incubated for 20 min at 37 °C to remove RNA. In order to amplify the SpCID1 gene from cDNA, gene specific primers CIDF and CIDR was used (Figure S1a, S2 and Table S1). Here as a positive control, GAPDH amplification was tested using primers GAPF and GAPR. For amplifying the gene, 2 μL of cDNA was incubated with forward and reverse primers (0.5 μM), dNTPs (0.2 mM), Phusion® High-Fidelity DNA Polymerase (0.1 U/μL) in 1X Phusion HF buffer in a final volume of 20 μL. PCR conditions: heat denaturation at 98 °C for 3 min, 35 cycles of (denaturing: 98 °C for 20 s, annealing: 58 °C for SpCID1 and 62.2 °C for GAPDH for 30 s, extension: 72 °C for 40 s), final extension at 72 °C for 5 min. The amplicons were visualized on 1.5% agarose gel run in 1X TAE buffer (Figure S1b). Further, the PCR product corresponding to SpCID1 gene was isolated using NucleoSpin® Gel and PCR Clean-up kit to obtain 2.65 μg of gene amplicon. To add overlapping regions corresponding to empty pET28a plasmid to the 3′ and 5′ of SpCID1 gene, vector complementary primers CIDMF and CIDMR (0.6 μM) were incubated with SpCID1 gene amplicon (100 ng), dNTPs (0.2 mM), PrimeSTAR GXL DNA polymerase (0.025 U/μL) in 1X PrimeSTAR GXL buffer in a final volume of 50 μL (Figure S1a, S2 and Table S1). PCR conditions: heat denaturation at 98 °C for 3 min, 35 cycles of (denaturing: 98 °C for 30 s, annealing: 55 °C for 30 s, extension: 68 °C for 1.33 min), final extension at 68 °C for 10 min. The product amplicon was isolated using Mega clear PCR clean-up kit to obtain 1.84 μg of 'megaprimer'.

In order to insert this 'megaprimer' into the bacterial expression vector, PCR was performed using 'megaprimer' (500 ng), pET28a plasmid (100 ng), dNTPs (0.2 mM), and PrimeSTAR GXL DNA polymerase (0.05 U/μL) in 1X PrimeSTAR GXL buffer in a final volume of 25 μL (Figure S1a, S2). PCR conditions: heat denaturation at 98 °C for 2 min, 35 cycles of (denaturing: 98 °C for 10 s, annealing: 65 °C for 30 s, extension: 68 °C for 6.5 min), final extension at 68 °C for 10 min. Further, digestion of unamplified pET28a plasmid was performed with Dpn1 (20 U) restriction enzyme, which specifically digests methylated recognition sites thereby leaving only the amplified clone in the PCR mixture. This reaction cocktail was transformed in chemically competent *E. coli* DH5α cells (by providing heat shock at 42 °C) and plated overnight at 37 °C on LB agar having kanamycin (50 μg/mL) to obtain bacterial colonies. In parallel, negative control was performed wherein, PCR and Dpn1 digestion was performed without addition of the 'megaprimer' followed by transformation, which gave no colonies. Plasmid from each colony was isolated using QIAprep Spin Miniprep kit using the manufacturer's protocol. In order to check for positive clones, isolated plasmid corresponding to DNA isolated from each colony was used as a template for PCR using T7 forward and reverse sequencing primer (T7F and T7R). The primers (0.5 μM), EmeraldAmp GT PCR master mix (10 μL), clone DNA (100 ng) was added to a final volume of 20 μL in autoclaved water. PCR conditions: heat denaturation at 94 °C for 3 min, 35 cycles of (denaturing: 94 °C for 30 s, annealing 58 °C for 30 s, extension: 68 °C for 1.5 min), final extension at 68 °C for 10 min. Also, as a positive control, PCR was performed with empty pET28a plasmid (100 ng). Further one of the positive clones was sequenced and was found to match with the desired pET28a_SpCID1 construct (Figure S3).

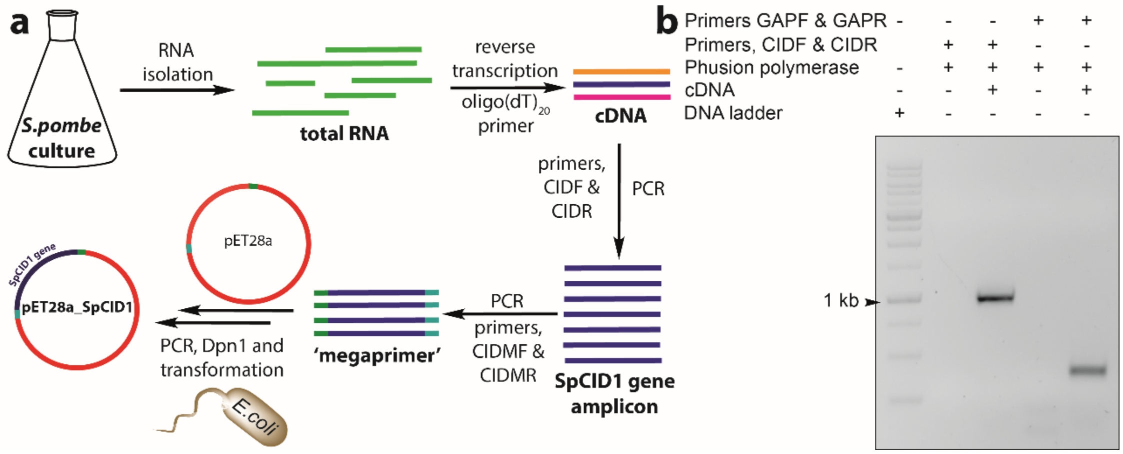


**Figure S1.** (**a**) Work-flow for isolating and inserting SpCID1 gene into pET28a plasmid. Total RNA from *S. pombe* was isolated and cDNA synthesized using oligo(dT)$_{20}$ primer. cDNA obtained was subjected to PCR using gene specific primers, CIDF and CIDR to obtain SpCID1gene amplicon. Further, overlapping regions corresponding to empty pET28a plasmid were added at 5′ and 3′ end using vector complementary primers CIDMF and CIDMR to obtain 'megaprimer'. Insertion of SpCID1 into pET28a plasmid was performed by PCR with 'megaprimer' followed by Dpn1 digestion. Transformation into chemical competent *E. coli* DH5α cells yielded the desired pET28a_SpCID1 plasmid. (**b**) Gene amplicons of SpCID1 (1014 bp) and GAPDH (400 bp) obtained upon PCR with cDNA was resolved on a 1.5% agarose gel and stained with SYBR Safe Stain.

5′...TCTCGATCCCGCGAAATTAATACGACTCACTATAGGGGAATTGTGAGCGGATAACAATTCCCCTCTAG
AAATAATTTTGTTTAACTTTAAGAAGGAGATATACCATGGGCAGCAGCCATCATCATCATCATCACAGCAG
CGGCCTGGTGCCGCGCGGCAGCCATATG**TCGCACAAGGAATTTACGAAGTTTTGCTATGAAGTGTAT**
**AATGAGATTAAAATTAGTGACAAAGAGTTTAAAGAAAAGAGAGCGGCATTAGATACACTTCGGCTA**
**TGCCTTAAACGAATATCCCCTGATGCTGAATTGGTAGCCTTTGGAAGTTTGGAATCTGGTTTAGCA**
**CTTAAAAATTCGGATATGGATTTGTGCGTGCTTATGGATTCGCGCGTCCAAAGTGATACAATTGCG**
**CTCCAATTCTATGAAGAGCTTATAGCTGAAGGATTTGAAGGAAAATTTTTACAAAGGGCAAGAATT**
**CCCATTATCAAATTAACATCTGATACGAAAAATGGATTTGGGGCTTCGTTTCAATGTGATATTGGAT**
**TTAACAATCGTCTAGCTATTCATAATACGCTTTTACTTTCTTCATATACAAAATTAGATGCTCGCCTA**
**AAACCCATGGTCCTTCTTGTTAAGCATTGGGCCAAACGGAAGCAAATCAACTCTCCTTACTTTGGA**
**ACTCTTTCCAGTTATGGTTACGTCCTAATGGTTCTTTACTATCTGATTCACGTTATCAAGCCTCCCG**
**TCTTTCCTAATTTACTGTTGTCACCTTTGAAACAAGAAAAGATAGTTGATGGATTTGACGTTGGTT**
**TTGACGATAAACTGGAAGATATCCCTCCTTCCCAAAATTATAGCTCATTGGGAAGTTTACTTCATG**
**GCTTTTTTAGATTTTATGCTTATAAGTTCGAGCCACGGGAAAAGGTAGTAACTTTTCGTAGACCAG**
**ACGGTTACCTCACAAAGCAAGAGAAAGGATGGACTTCAGCTACTGAACACACTGGATCGGCTGAT**
**CAAATTATAAAAGACAGGTATATTCTTGCGATTGAAGATCCTTTCGAGATTTCACATAATGTGGGTA**
**GGACAGTTAGCAGTTCTGGATTGTATCGGATTCGAGGGGAATTTATGGCCGCTTCAAGGTTGCTC**
**AATTCTCGCTCATATCCTATCCCTTATGATTCATTATTTGAGGAGGCC**TAGCTCGAGCACCACCACCA
CCACCACTGAGATCCGGCTGCTAACAAAGCCCGAAAGGAAGCTGAGTTGGCTGCTGCCACCGCTGAGC
AATAACTAGCATAACCCCTTGGGGCCTCTAAAC...3′

**Figure S2.** Sequence of SpCID1 gene in desired pET28a_SpCID1 plasmid (bold) and sequence corresponding to gene specific primers used for getting gene amplicon (blue). Sequence corresponding to vector complementary primer used for generating 'megaprimer' (underlined red). Sequencing primers T7 forward and reverse are underlined in black (See Table S1).

**Table S1.** Sequence of DNA primers and templates used.

| Primers and Templates | Sequence |
|---|---|
| sgRNA forward primer, **CFP1** | 5' GCGTAATACGACTCACTATAGGGTTAGG 3' |
| sgRNA forward primer, **CFP1′** | 5' GCGTAATACGACTCACTATAGTTAGGGTTAG 3' |
| sgRNA forward primer, **CFP2** | 5' GCGTAATACGACTCACTATAGGCGA 3' |
| sgRNA reverse primer, **CRP** | 5' TTTTTTTTTTTTTTTAAAGCACCGACTCGGTGC 3' |
| sgRNA template, **CT1** | 5' GCGTAATACGACTCACTATAGGGTTAGGGTTAGGGTTAGGGTTAGTTTAAGAGCTA TGCTGGAAACAGCATAGCAAGTTTAAATAAGGCTAGTCCGTTATCAACTTGAAAAAG TGGCACCGAGTCGGTGCTTT 3' |
| sgRNA template, **CT1′** | 5' GCGTAATACGACTCACTATAGTTAGGGTTAGGGTTAGGGTTGTTTAAGAGCTATGC TGGAAACAGCATAGCAAGTTTAAATAAGGCTAGTCCGTTATCAACTTGAAAAAGTGG CACCGAGTCGGTGCTTT 3' |
| sgRNA template, **CT2** | 5' GCGTAATACGACTCACTATAGGCGAGGGCGATGCCACCTAGTTTAAGAGCTATGC TGGAAACAGCATAGCAAGTTTAAATAAGGCTAGTCCGTTATCAACTTGAAAAAGTGG CACCGAGTCGGTGCTTT 3' |
| eGFP template forward primer, **eGFPF** | 5' AGGGCGAGGAGCTGTTCA 3' |
| eGFP Template reverse primer, **eGFPR** | 5' GGTAGCGGCTGAAGCACT 3' |
| qPCR telomere forward primer, **TeloF** | 5' GGTTTTTGAGGGTGAGGGTGAGGGTGAGGGTGAGGGT 3' |
| qPCR telomere reverse primer, **TeloR** | 5' TCCCGACTATCCCTATCCCTATCCCTATCCCTATCC-CTA 3' |
| 5′ FAM-labeled target DNA strand 1, **TS1** | 5' FAM TAATGAATTCCCCAATACCCTAACCCTAACCCTAACCCTAACCCGTTCATAT AA 3' |
| Target DNA strand 2, **TS2** | 5' TTATATGAACGGGTTAGGGTTAGGGTTAGGGTTAGGGTATTGGGGAATTCATTA 3' |
| Gene specific forward primer, **CIDF** | 5' TCGCACAAGGAATTTACGAAGTTTTGC 3' |
| Gene specific reverse primer, **CIDR** | 5' GGCCTCCTCAAATAATGAATCATAAGGG 3' |
| GAPDH forward primer, **GAPF** | 5' TGGCAATTCCTAAGGTTGGTATTAACGGTT 3' |
| GAPDH reverse primer, **GAPR** | 5' CCGACAACGTACATGGGGGCGTC 3' |
| Vector complementary forward primer, **CIDMF** | 5' GGTGCCGCGCGGCAGCCATATGTCGCACAAGGAATTTACGAAGTTTTGC 3' |
| Vector complementary reverse primer, **CIDMR** | 5' CAGTGGTGGTGGTGGTGGTGCTCGAGCTAGGCCTCCTCAAATAATGAATCATAAG GG 3' |
| Sequencing primer T7 forward, **T7F** | 5' CGCGAAATTAATACGACTCACTATAGGG 3' |
| Sequencing primer T7 reverse, **T7R** | 5' GTTATGCTAGTTATTGCTCAGCGG 3' |

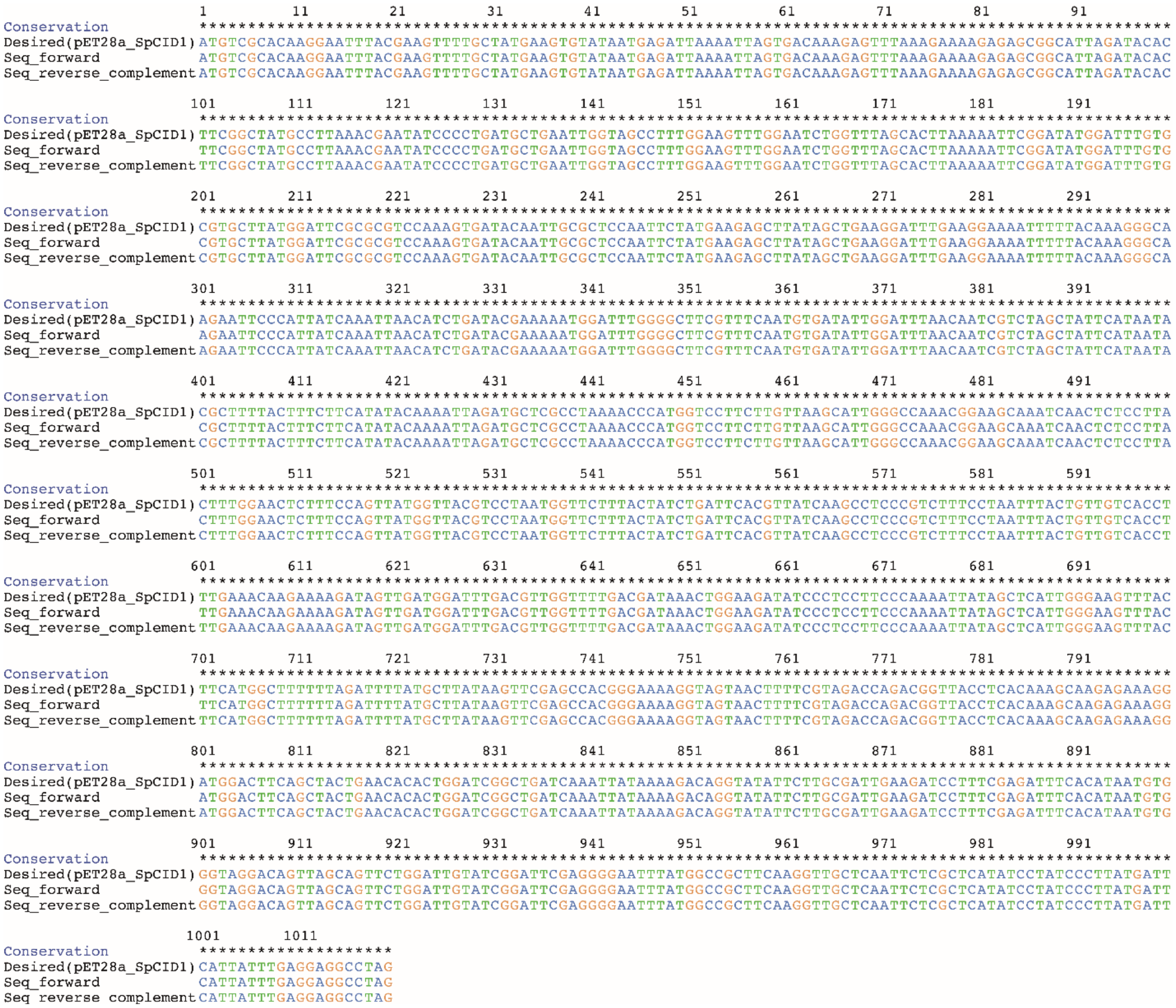

**Figure S3.** Alignment of SpCID1 gene sequence in the desired plasmid (pET28a_SpCID1) with forward and reverse complement of sequencing results (obtained with sequencing primers T7F and T7R, Table S1). Sequence conservation denoted by Asterix. Alignment of sequencing results performed using ClustalW and represented by UCSF Chimera software.

**SpCID1 expression and purification**

The plasmid, pET28a_SpCID1 was transformed into chemically competent Rosetta™(DE3) cells and streaked on LB agar plate having both kanamycin (50 µg/mL) and chloramphenicol (25 µg/mL). After 16 h, a colony was picked from the plate and incubated in primary culture in LB with both the antibiotics until $OD_{600}$ of ~1.4 was reached. Two percent of the primary culture was inoculated in secondary culture in multiple batches of 1 L volume LB cultures having both the antibiotics. Further, the cultures were induced with 0.5 mM IPTG at $OD_{600}$ of ~0.6 and grown at 18 °C for 16 h with 180 rpm shaking. The cells were then pelleted down and resuspended in 50 mL lysis buffer (50 mM Tris pH 7.5, 150 mM NaCl, 10 mM Imidazole, 30 µM PMSF), sonicated (20 s pulse on, 30 s pulse off at 80% amplitude) for 10 min and the lysate obtained was centrifuged at 14,500 rpm for 30 min. Further to purify the protein by immobilized metal affinity chromatography (IMAC), the supernatant was incubated with 10 mL of Ni-IDA resin for 90 min under slow rotation at 4 °C and the resin was packed in a glass column. The resin was washed with 500 mL of wash buffer (50 mM Tris pH 7.5, 150 mM NaCl, 75 mM imidazole, 30 µM PMSF) and protein was eluted in 25 mL elution buffer (50 mM Tris pH 7.5, 150 mM NaCl, 250 mM imidazole, 30 µM PMSF). The eluted protein was buffer exchanged with size exclusion buffer (20 mM HEPES pH 6.8, 150 mM NaCl, 5 mM 2-mercaptoethanol, 30 µM PMSF) at 4 °C and concentrated using Vivaspin columns.  The concentrated enzyme (10 mL) was purified by FPLC on HiLoad 16/600 Superdex 200 preparative column and eluted in size exclusion buffer at a flow rate of 1 mL/min. Fractions corresponding to SpCID1 enzyme were dialysed three times against protein storage buffer (20 mM HEPES pH 6.8, 150 mM NaCl, 5 mM 2-mercaptoethanol, 50% glycerol), aliquoted and stored at -40 °C. The protein concentration was calculated using Bradford assay against a standard solution of BSA, which yielded a protein stock of 10.25 µM (Figure S4). Molecular weight of SpCID1 is 40,968 Da.

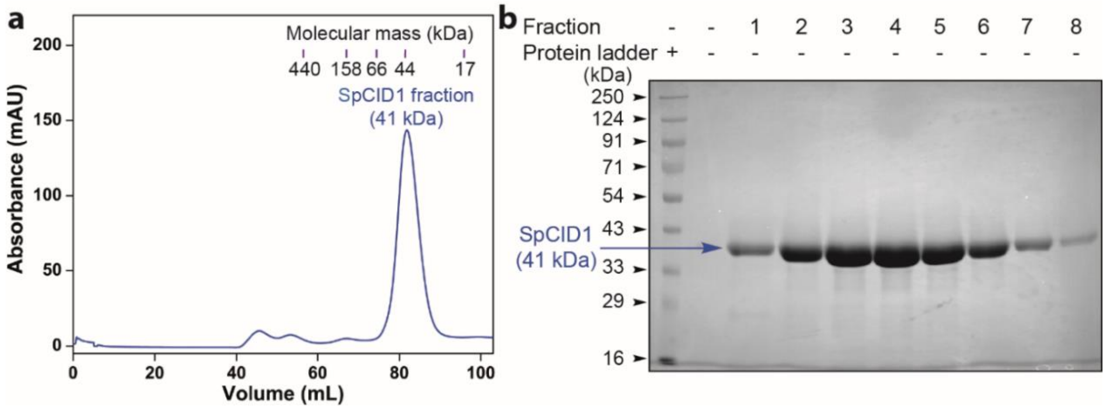


**Figure S4.** (**a**) Size exclusion chromatogram of IMAC purified SpCID1 enzyme (~41 kDa) as compared with molecular mass of protein standards. (**b**) Fractions corresponding to SpCID1 were loaded and resolved by 12.5% SDS-PAGE and stained with Coomassie blue.

**Conditions for single nucleotide incorporation**

Terminal uridylation at the 3′ end of a model RNA oligonucleotide was optimized by incubating 5′-FAM-labeled RNA (10 μM) with AMUTP (10 μM), APUTP (25 μM) or ATUTP (50 μM) in the presence of Tris-HCl buffer (10 mM, pH 7.9 at 25 °C), NaCl (50 mM), $MgCl_2$ (10 mM), DTT (2 mM), RiboLock RNase inhibitor (1 U/μL) and SpCID1 (3.42 pmol for AMUTP and APUTP, 10.25 pmol for ATUTP) in a final volume of 20 μL. After 5, 15 and 30 mins, 5 μL reaction aliquots were mixed with denaturing loading buffer (15 μL) and heat-denatured at 75 °C for 3 min. 5 μL of the sample was loaded on to 20% denaturing polyacrylamide gel, electrophoresed and imaged using Typhoon gel scanner at FAM wavelength.

***Large-scale synthesis of single azide-modified RNAs:*** 5′-FAM-labeled RNA oligonucleotide (10 μM, 200 pmol) was incubated with AMUTP (10 μM), APUTP (25 μM) or ATUTP (50 μM) in the presence of Tris-HCl buffer (10 mM, pH 7.9 at 25 °C), NaCl (50 mM), $MgCl_2$ (10 mM), DTT (2 mM), RiboLock RNase inhibitor (1 U/μL) and SpCID1 (3.42 pmol for AMUTP and APUTP, 10.25 pmol for ATUTP) in a final volume of 20 μL and was incubated for 15 min in case of AMUTP and APUTP or 30 min in case of ATUTP. Multiple such reactions (e.g., 5 sets) were performed and heat-inactivated at 75 °C for 3 min. Reaction sets corresponding to an individual nucleotide were combined and purified by RP-HPLC. A 5 set reaction (1 nmol) yielded 89%, 95% and 76% of the 3′-AMU, 3′-APU or 3′-ATU-labeled RNA products. ESI-MS analysis was performed in negative mode by injecting RNA oligonucleotide products (100 pmol) dissolved in 50% acetonitrile in an aqueous solution of 10 mM triethylamine and 100 mM hexafluoro-2-propanol See Table S2 for yield and mass data.

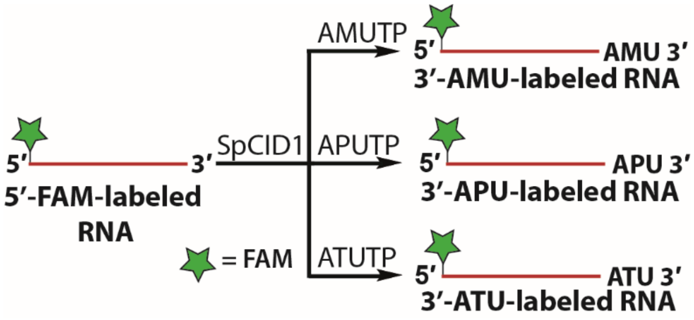


**Figure S5.** Single nucleotide incorporation of azide-modified UTPs (AMUTP, APUTP and ATUTP) at the 3′ end of 5′-FAM-labeled RNA oligonucleotide using SpCID1.

**Table S2.** Mass and yield data of RNA oligonucleotide products of terminal uridylation and click reactions.

| RNA oligonucleotide product | calcd. mass | found mass | isolated yield (nmol) | isolated yield (%) |
|---|---|---|---|---|
| 3′-AMU-labeled RNA | 9750.0 | 9749.6 | 0.89[a] | 89 |
| 3′-APU-labeled RNA | 9778.1 | 9777.6 | 0.95[a] | 95 |
| 3′-ATU-labeled RNA | 9950.3 | 9949.8 | 0.76[a] | 76 |
| Cy3-labeled RNA | 10733.2 | 10732.6 | 0.15[b] | 51 |
| Alexa 594-labeled RNA | 10509.9 | 10509.1 | 0.16[b] | 52 |

Isolated yields are with respect to [a]1 and [b]0.3 nmol of the substrate RNA oligonucleotide.
See Figure S6 for mass spectra.

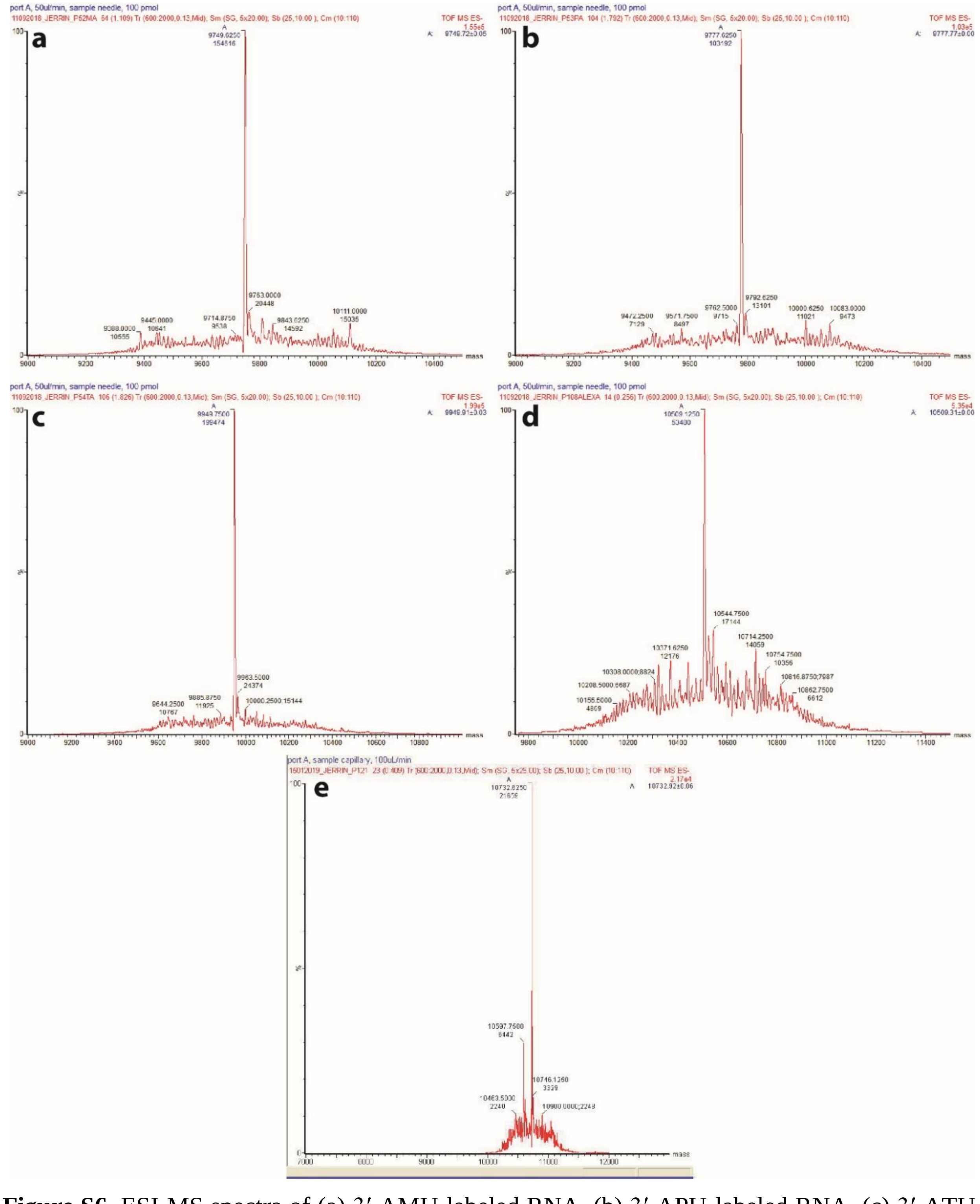

**Figure S6.** ESI-MS spectra of (a) 3′-AMU-labeled RNA, (b) 3′-APU-labeled RNA, (c) 3′-ATU-labeled RNA, (d) Alexa-labeled RNA and (e) Cy3-labeled RNA (see Table S2 for details).

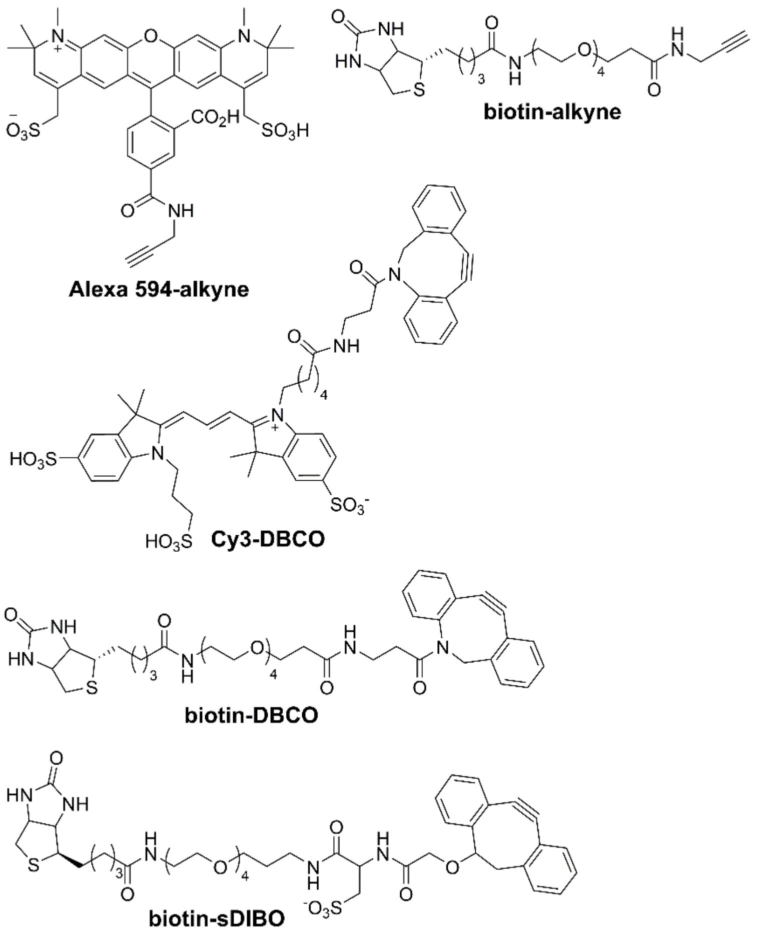


**Figure S7.** Structure of terminal and strained alkyne substrates used in CuAAC and SPAAC reactions.

### Click reaction on model 3′-AMU-labeled RNA oligonucleotide

***Strain-promoted azide-alkyne cycloaddition (SPAAC) reaction:*** 3′-AMU-labeled RNA oligonucleotide (10 μM, 300 pmol) was incubated with Cy3-DBCO (1 mM), in a final volume of 30 μL in autoclaved water (Figure S7). The reaction was incubated at 37 °C for 2 h followed by addition of 30 μL of 5 M ammonium acetate and 300 μL ethanol. The reaction mixture was precipitated at -20 °C overnight followed by centrifugation at 15,000 rpm for 15 min. The pellet was washed with 500 μL pre-chilled 75% ethanol in autoclaved water to remove salts and unconjugated dye followed by centrifugation (15,000 rpm) for 15 min. The clicked RNA product was (Cy3-labeled RNA) dissolved in autoclaved water. See Table S2 for yield and mass data.

***Copper-catalyzed azide-alkyne cycloaddition (CUAAC) reaction:*** The catalyst mix was prepared by reducing $CuSO_4$ (0.2 mM) using sodium ascorbate (2 mM) in the presence of Cu(I)-stabilizing ligand THPTA (1 mM). To above solution was added Alexa 594-alkyne (1 mM, Figure S7) and

3′-AMU-labeled RNA oligonucleotide (10 µM, 300 pmol). The final reaction volume was adjusted to 30 µL and incubated at 37 °C for 2 h. To the reaction mix was added 30 µL 5 M sodium acetate and 300 µL ethanol and stored at -20 °C overnight. The sample was centrifuged at 15,000 rpm for 15 min and the pellet was washed with 500 µL of pre-chilled 75% ethanol in autoclaved water. The sample was again centrifuged at 15,000 rpm for 15 min and Alexa 594-labeled RNA product was dissolved in autoclaved water. See Table S2 for yield and mass data.

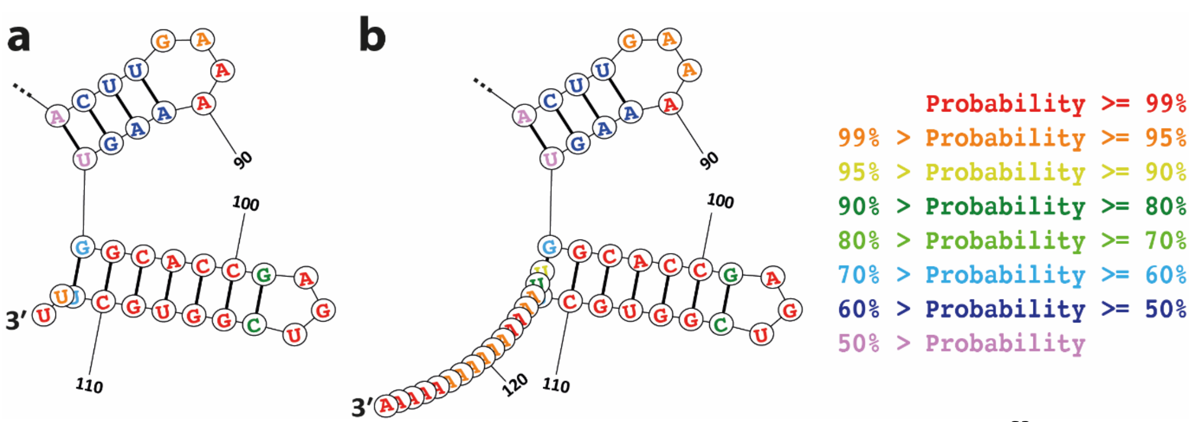


**Figure S8.** Structure prediction of sgRNAs using online software, RNAstructure.[S3] Structure of 3′ end of (**a**) conventional sgRNA design[S4] and (**b**) 15 adenylate tailed sgRNA is shown here. Color code for probability of structure formation is also given.

**Table S3.** Yield data for the synthesis of AMU-labeled sgRNAs and click-functionalized sgRNAs.

| AMU- and click-labeled sgRNAs | isolated yield (nmol) | isolated yield (%) |
|---|---|---|
| sgRNA $\mathbf{1}_{\mathbf{Az}}$ | 0.90[a] | 90 |
| sgRNA $\mathbf{1'}_{\mathbf{Az}}$ | 1.86[b] | 93 |
| sgRNA $\mathbf{2}_{\mathbf{Az}}$ | 1.91[b] | 96 |
| sgRNA $\mathbf{1}_{\mathbf{Cy3}}$ | 0.27[c] | 90 |
| sgRNA $\mathbf{1'}_{\mathbf{Cy3}}$ | 1.66[d] | 92 |
| sgRNA $\mathbf{2}_{\mathbf{Cy3}}$ | 1.65[d] | 92 |
| sgRNA$\mathbf{1}_{\mathbf{Bio}}$ | 0.23[c] | 77 |

Isolated yields are with respect to [a]1, [b]2, [c]0.3, [d]1.8 nmol of the substrate RNA.

**Functionalization of azide-labeled sgRNAs using click chemistry**

In order to conjugate azide-labeled sgRNA with various functional tags, sgRNA $\mathbf{1_{Az}}$, $\mathbf{1'_{Az}}$ or $\mathbf{2_{Az}}$ (10 μM) was incubated with Cy3-DBCO or biotin-DBCO (1 mM) in a final volume of 30 μL for 2 h at 37 °C (Figure S7). The products obtained were precipitated overnight using 1 volume of 5 M ammonium acetate and 10 volumes of ethanol and centrifuged at 15000 rpm for 15 min. RNA pellet obtained was washed with 75% ethanol in water and then centrifuged to afford the clicked products. Reacting $\mathbf{1_{Az}}$, $\mathbf{1'_{Az}}$ and $\mathbf{2_{Az}}$ with Cy3-DBCO yielded $\mathbf{1_{Cy3}}$, $\mathbf{1'_{Cy3}}$ and $\mathbf{2_{Cy3}}$, respectively. Similarly, reacting $\mathbf{1_{Az}}$ with biotin-DBCO yielded $\mathbf{1_{Bio}}$. The purified products were analysed by PAGE (8.5%) under denaturing conditions and imaged in Typhoon gel scanner in Cy3 wavelength and or stained with Stains-All reagent. See Table S2 for yields.

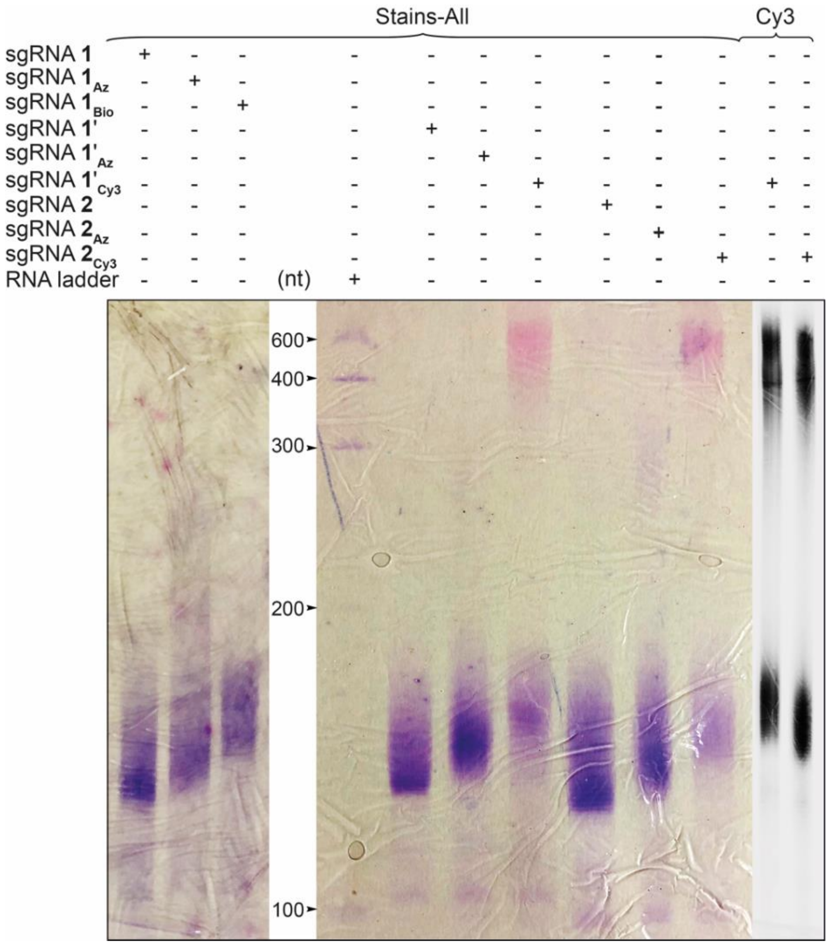


**Figure S9.** Terminal uridylation products $\mathbf{1_{Az}}$, $\mathbf{1'_{Az}}$, $\mathbf{2_{Az}}$ and product of SPAAC reaction with Biotin-DBCO and Cy3-DBCO, sgRNA $\mathbf{1_{Bio}}$, $\mathbf{1'_{Cy3}}$, $\mathbf{2_{Cy3}}$ were resolved and visualized by 8.5% denaturing PAGE using Typhoon gel scanner at Cy3 wavelength and or using Stains-All reagent.

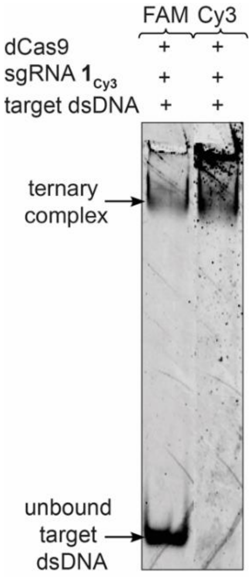


**Figure S10.** The ternary complex of Cy3-labeled sgRNA $\mathbf{1_{Cy3}}$, dCas9 and target telomere dsDNA visualized in FAM and Cy3 wavelength in gel scanner. The decreased binding of sgRNA $\mathbf{1_{Cy3}}$ is evident from the higher amount of unbound telomere dsDNA in FAM wavelength.
